# Pseudotime analysis reveals novel regulatory factors for multigenic onset and monogenic transition of odorant receptor expression

**DOI:** 10.1101/2022.01.31.478392

**Authors:** Mohammad Hussainy, Sigrun I. Korsching, Achim Tresch

**Author notes:** These authors contributed equally to this work: Sigrun I. Korsching; Achim Tresch.

## Abstract

During their maturation from horizontal basal stem cells, olfactory sensory neurons (OSNs) are known to select exactly one out of hundreds of olfactory receptors (ORs) and express it on their surface, a process called monogenic selection. Monogenic expression is preceded by a multigenic phase during which several OR genes are expressed in a single OSN. Here, we perform pseudotime analysis of a single cell RNA-Seq dataset of murine olfactory epithelium to precisely align the multigenic and monogenic expression phases with the cell types occurring during OSN differentiation. In combination with motif analysis of OR gene cluster-associated enhancer regions, we identify known and novel transcription (co-)factors (Ebf1, Lhx2, Ldb1, Fos and Sspp2) and chromatin remodelers (Kdm1a, Eed and Zmynd8) associated with OR expression. The inferred temporal order of their activity suggests novel mechanisms contributing to multigenic OR expression and monogenic selection.

## Introduction

The sense of smell is tasked with the daunting challenge of making sense of potentially billions of different chemical stimuli to enable a multitude of different behaviors such as food search, prey hunting, predator evasion, mating and other social interactions^1^. This task is solved by several different receptor families, of which the best studied is both the evolutionary oldest and the largest (odorant receptor genes, OR genes)^2^. The analytical power of this system is maximal when information gathered from activation of individual receptors is kept separate at the peripheral level. Indeed for both vertebrates and insects it has been shown that individual olfactory sensory neurons (OSNs) express only a single OR gene out of the entire olfactory receptor repertoire, which has been christened as monogenic expression^3,4^. Sensory neurons expressing the same receptor are distributed across the olfactory sensory surface, but their axons converge into a single target region in the olfactory bulb, the first relay station of olfactory information processing^5–8^. These target regions (so-called glomeruli) show a stereotyped arrangement, resulting in a receptotopic map on the olfactory bulb (or antennal lobe in the case of insects). Thus monogenic expression has a central importance for the olfactory coding logic. In fact, expression of ORs is even monoallelic, i.e. restricted to one allele of the OR selected in monogenic expression^3,4,9,10^. The molecular path towards monogenic and monoallelic expression is still not well understood, and the relative timing of these processes is not clear.

To reach monogenic and monoallelic expression presents a massive challenge in the case of very large gene families such as those of mouse and rat ORs, which both number well over one thousand intact genes^2^. A striking feature of genomic arrangement of OR genes is the occurrence in several clusters, which contain from a single to over one hundred different OR genes^11^. For mouse 68 such clusters have been identified, with the largest cluster containing 269 OR genes^11^. Another large cluster contains all 145 class I OR genes, which show a spatially restricted expression pattern in the olfactory epithelium^12–14^. These observations have prompted the search for cluster-specific regulatory elements. In a seminal publication, 63 genomic regions containing such elements were identified and named after Greek islands^11,15^. Fourty-two class II OR clusters and the single class I OR cluster are associated with these Greek islands, which lie proximal to and sometimes even inside the clusters^11^. A common feature of Greek islands is the presence of closely adjacent Lhx2 and Ebf1-binding motifs, which are also found individually in promoter regions of individual OR genes^16–19^.

Beyond individual cluster-specific regulatory elements the chromatin structure itself appears to play an essential role in regulating OR expression. OSNs possess a unique nuclear architecture compared to other cell types including the basal cells giving rise to the OSN lineage. In the silent phase before onset of expression OR genes are aggregated in constitutive heterochromatin and are associated with its molecular hallmarks, H3K9me3 and H4K20me3^20,21^. Onset of expression is concomitant with selective de-methylation (H3K9me3), methylation of H3K27me3 and re-location into expression-competent territory^21–24^. Moreover, of the two alleles of an active OR only one is found in the more plastic facultative heterochromatin^22–24^, i.e. amenable to expression, whereas the other remains blocked inside the constitutive heterochromatin, resulting in monoallelic expression. This suggests an involvement of chromatin remodelers in regulation of expression of OR genes. Furthermore, the stabilization of monogenic expression appears to require negative feedback from an active OR gene^25,26^, which may be mediated by silencing of the activating demethylase LSD1, synonym Kdm1a^27,28^.Recent progress in deep sequencing techniques has allowed to obtain high quality single cell transcriptomes (scRNA-Seq), resulting in the surprising observation that monogenic expression of ORs found in mature OSNs is preceded by a multigenic phase in immature OSN^29,30^.

It is so far mostly unclear how these OR expression phases align to the developmental stages of OSN differentiation. Furthermore, although the basic stages (stem cell, dividing precursor cell, immature neuron, mature neuron) were known previously^29,30^, deep sequencing techniques allow an unbiased ordering of individual cells along pseudotime trajectories according to their entire transcriptome. This enables a more precise and more stringent categorization of developmental stages compared to previous attempts.

Here, we re-analyzed a scRNA-Seq dataset obtained by Fletcher et al^31^ with the goal to precisely determine the timing of multigenic and monogenic expression during OSN differentiation. A combination of sequence binding motif and time series analysis then identifies novel regulatory components involved in establishing OR gene expression patterns. We ascertain the transcription (co)factors and chromatin remodelers that are specifically correlated with the onset of multigenic and of monogenic expression (e.g., Fos, Ssbp2, Eed and Zmynd8). Finally, we suggest potential mechanisms for multigenic and monogenic selection.

## Results

### Re-analysis of a single cell RNA-Seq data set reveals four lineages originating from globose basal cells

We re-analyzed a scRNA-Seq dataset obtained from Fletcher et al^31^. After quality filtering and pre-processing (Methods, SFigure 2,3), 687 cells were included into further analysis and grouped into 13 clusters using Seurat KNN clustering on the top 15 principal components (Methods). Dimension reduction and visualization was performed using principal components analysis (Figure 1A) and tSNE / UMAP. Using an extensive set of known marker genes, we assigned clusters to cell types of the main olfactory epithelium (MOE) (Figure 1B, Methods SFigure 4A). We detected all cell types described by Fletcher et al^31^, and additionally we could subdivide the globose basal stem cell cluster into quiescent cells (qGBC) and active cells (GBC). Active GBC were identified by their expression of Ascl1/Mash1^32–35^ and by the presence of cell cycle genes such as Mki67 and Top2a. GBC are known as the adult OSN stem cells responsible for a sustained self-renewal of the OSNs throughout life^35^.

**Figure 1.**
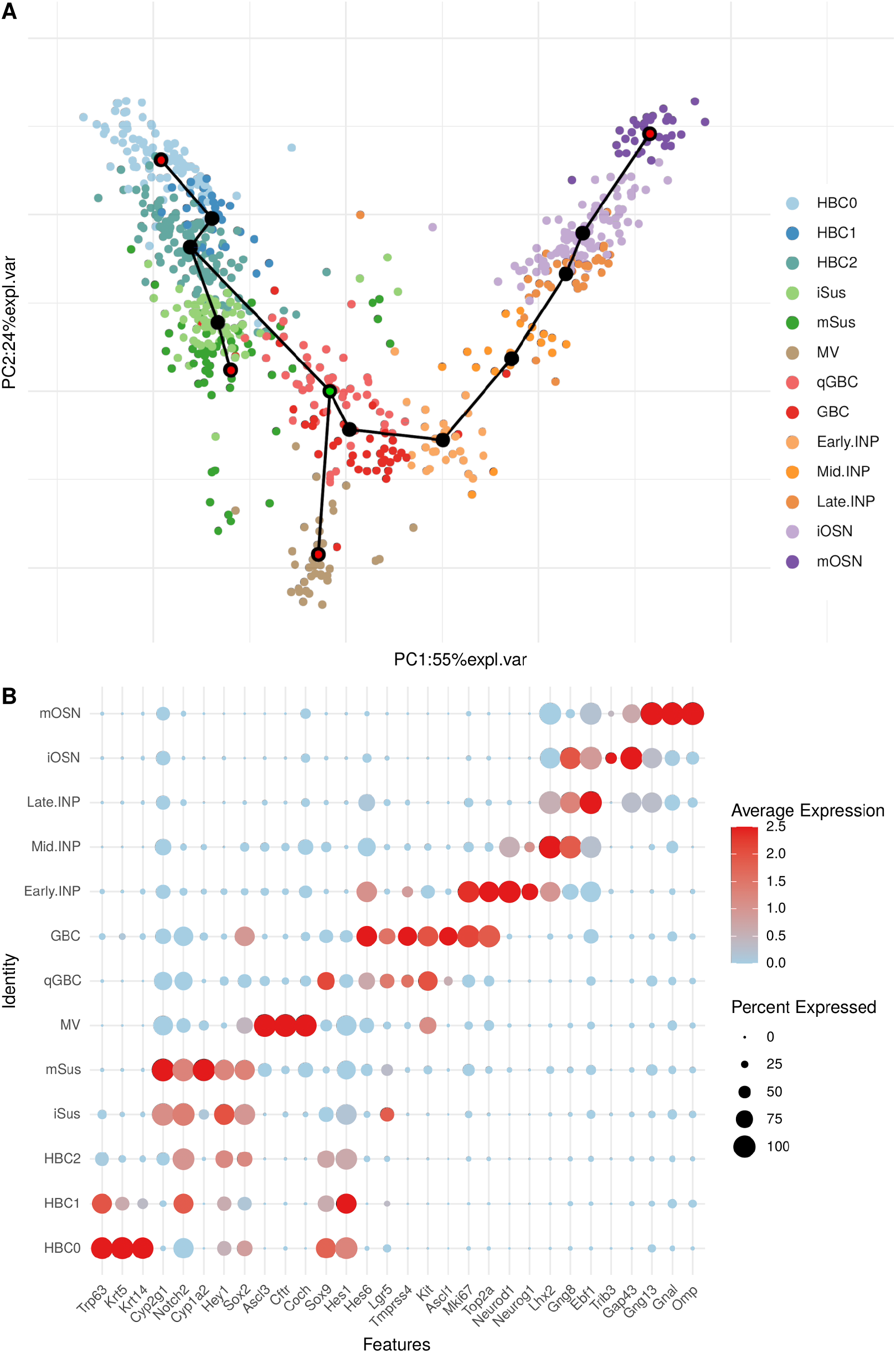
Cell type identification, trajectory inference and pseudotime assignment. **A)** 2D PCA projection of MOE cells shows the four predicted lineages starting from qGBC (center, pink, the starting point of the four trajectories is marked in green). The differentiation trajectory end points (red dots at the end of paths) are HBC0 (top left, pale blue), mSus (center left, green), MV (bottom, brown) and mOSN (top right, purple). The transition stages in the four lineages (black dots) are HBC1 (light blue), HBC2 (blue), iSus (light green), GBC (red), Early.INP (pale orange), Mid.INP (light orange), Late.INP (dark orange) and iOSN (light purple) from the left to the right. **B)** DotPlot representation of the average expression (red for high expression and pale blue for lower one) of known marker genes (x.axis) corresponding to cell types of MOE (y.axis), and the size of each dot represents the percentage of cells that expresses a corresponding marker gene in a given cell type.

Further, trajectory inference by Slingshot^36^ (Methods) revealed a tree with four leaves (Figure 1A). Be aware that Slingshot does not provide information on the direction of development, but merely a tree topology. In the following, we restrict our analysis to the neuronal lineage and therefore choose qGBC as a root node for pseudotime analysis. qGBC has been reported as a general stem cell population in the MOE following injury^35^. A previous analysis by Fletcher et al^31^ focused on differentiation processes starting from HBC0. They placed their root node in HBC0, which is a leaf node of our tree, and therefore merely report three lineages. Apart from this, their reconstruction is essentially identical to ours. The four branches of our tree are (Figure 1A):

1. The basal stem cells lineage, which connects the quiescent horizontal basal stem cells (HBC0, represented by 91 cells in the data) via two transient populations of horizontal basal stem cells (HBC1 (29 cells) and HBC2 (118 cells)) with the qGBC (53 cells).
2. The supporting cells lineage ranges from mature sustentacular cells (mSus (70 cells)) to qGBCs, and includes immature sustentacular cells (iSus (49 cells)) and HBC2.
3. The microvillous cells lineage contains merely microvillous cells (MV (36 cells)). No transient cell types have been detected in this trajectory.
4. The neuronal lineage ends with mature olfactory sensory neurons (mOSN (32 cells)). It spans a range of several stages, namely GBC (34 cells), three intermediate neuronal precursors Early.INP (26 cells), Mid.INP (20 cells), Late.INP (34 cells) and immature olfactory sensory neurons (iOSN (95 cells)).

### OR gene expression is limited to the last three stages of OSN differentiation: Sudden onset of multigenic OR expression in Late.INP is followed by transition to monogenic expression in immature OSN stage

Expectedly, OR expression is essentially unique to the neuronal lineage (Figure 2). Next we used Slingshot to assign a pseudotime to each cell, thereby providing a linear order of all 294 cells in the neuronal lineage from qGBC to the terminal cell cluster (Methods). Our analysis could detect 157 cells of neuronal lineage that express at least one OR gene at relevant levels, i.e. ≥50 normalized counts. 132 of them belonged to the last three stages of OSN differentiation. Most of Late.INP cells (28 of 34), iOSN (76 of 95) and mOSN (28 of 32) express at least one OR. Many cells express more than one OR (multigenic expression), in particular in the Late.INP stage (26 of 28 cells expressing OR genes, 92.8%). The frequency of multigenic expression drops sharply in later stages, 42% and 32% for iOSN and mOSN, respectively. We found 212 different OR genes that were expressed at least once in a single cell of the neuronal lineage (Excel file 1). Figure 2A shows the total number of reads that were assigned to OR genes, separately for each cell. While aggregate OR expression levels are almost zero for qGBC/GBC, early and mid INPs, there is a steep onset of OR expression in Late.INP. Then, overall OR expression stays at similarly high levels in iOSN and mOSN (Figure 2A). A few OSN do not appear to express any OR at a relevant level. More precisely,19 out 95 iOSN cells (20%) and 4 out 32 mOSN (12.5%) have less than 50 normalized OR counts. While it cannot be excluded that this is caused by incomplete annotation of the OR repertoire, it is also possible that reads were excluded due to multiple mapping to closely similar OR genes.

**Figure 2.**
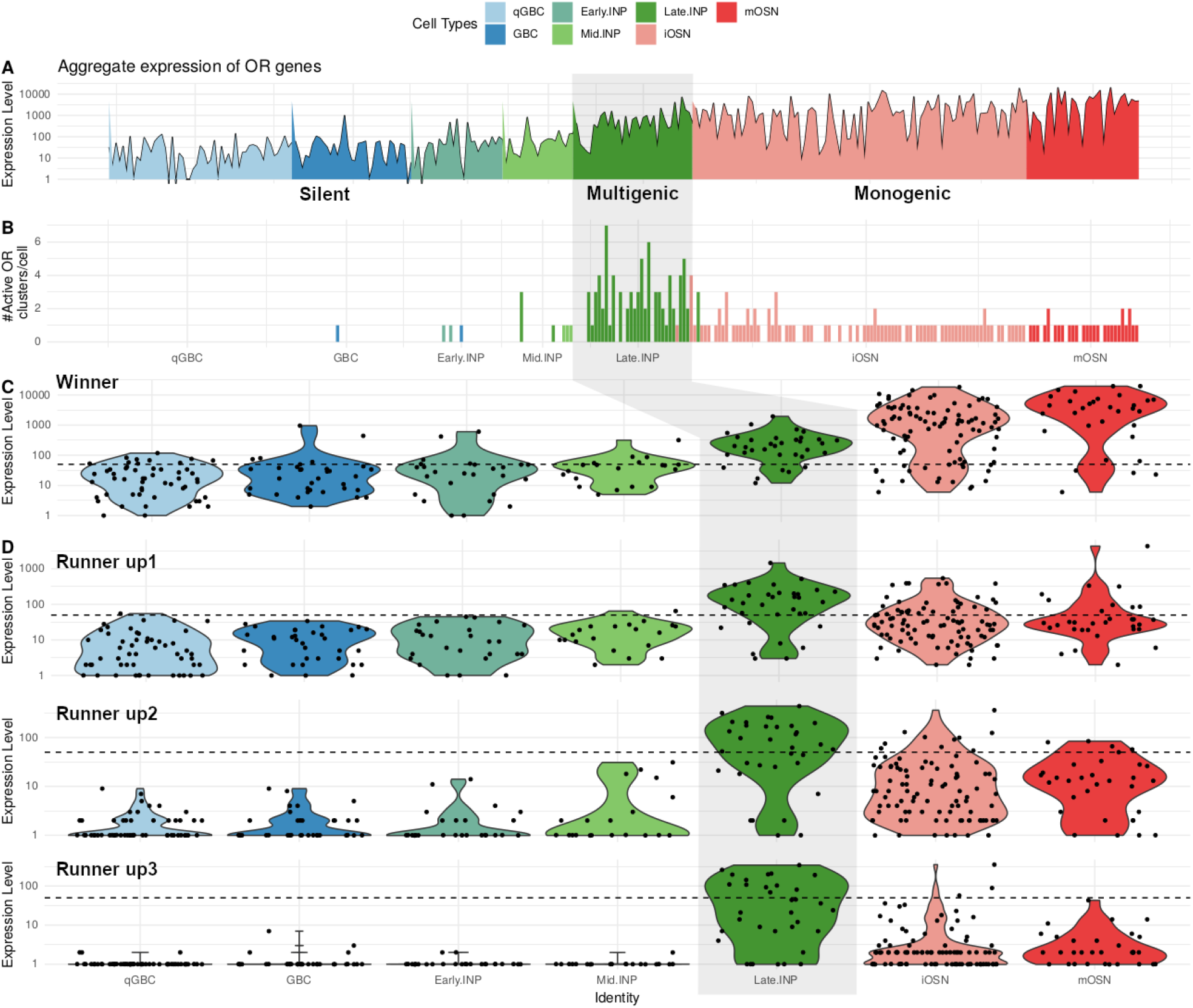
OR expression dynamics along neuronal lineage. **A)** Aggregate expression (normalized pseudocounts) of all OR genes for each cell. Cells are sorted according to pseudotime and colored according to cell type. **B)** Number of active OR clusters per cell, sorted by pseudotime. **C)** Expression of the OR gene with the highest expression level in each cell (“winner”). For each stage of the neuronal lineage, we show the distribution of the corresponding expression values (pseudocounts) as a violin plot. **D)** Using the same representation as in C, the expression of the OR gene with second, third and fourth highest expression (“runners up” 1-3) in each cell is shown in the top, middle, and bottom row, respectively. Note that the y-axis is in log scale, and that the scale of the winner expression is on average one to two orders of magnitude higher than that of the runners up. The dotted horizontal line in **C)** and **D)** marks an expression level of 50 counts.

It is known that mature OSNs express only a single OR gene^3,4,9,37^, after a transient period of multigene expression^29,30^. We therefore decided to rank OR genes by expression level, separately in each cell. We then investigated the temporal behavior of the top four ranked OR genes in each cell. These genes account for 99% of all reads (413,388 out of 416,033) mapped to OR genes in cells from the neuronal lineage. The top-ranked gene of each stage will be referred to as ‘winner’ and the others as the ‘runners-up’. While the abundance of all runners-up drops sharply after Late.INP, the winner does not drop and in fact absolute levels keep increasing several fold until the mOSN stage (Figure 2C,D). As a consequence the distance between winner and runners-up increases considerably in the iOSN stage and even more so in the mature neurons (mOSN). Since we observe each cell only once, we cannot be sure that the winner gene observed in one cell at a certain stage will be the highest expressed OR gene when that cell matures. However, this is by far the most plausible explanation, since a rank switch between the winner and a runner-up before / during iOSN stage would require a coordinated switch of expression between these two specific OR genes, from high to almost zero and vice versa, an unlikely scenario.

Taken together, the pseudotime analysis of neuronal lineage cells suggests three main phases for OR expression (Figure 2C,D):

1. The silent phase exhibits virtually no OR expression, which is the case for the four early stages of neuronal lineage (qGBC, GBC, Early.INP and Mid.INP). In the present data, the silent phase is represented by 133 cells.
2. The onset of OR expression (multigenic phase) is characterized by the simultaneous expression of several OR genes per cell at relatively similar levels: this phase is represented by 34 cells and contains specifically the Late.INP stage and the beginning of the iOSN stage.
3. Finally, the monogenic phase includes the end of the iOSN stage and the mOSN stage, where each cell expresses essentially one functional OR allele (henceforth called the “winner”, Figure 2C) at a very high level while the remaining ORs (the “runners-up”, Figure 2D) show no or very low expression. Our data contains 127 cells in this phase.

Our analysis reveals that Late.INP is a crucial stage in stochastic selection.

### OR gene expression during multigenic phase shows no sign of OR cluster-specific activation

Next, in an attempt to infer the mechanism of activation in the multigenic stage and transition to monogenic stage, we analyzed the joint OR gene expression per cell (Figure 2B, Excel file 2). We found that each of the cells expressing more than one OR gene in the Late.INP stage had at least two active OR clusters (i.e., clusters with an OR gene expressed at a level of at least 50 counts). The number of active clusters reached up to 7 for some cells. We performed a permutation test to assess whether OR genes that are jointly active in one cell have the tendency to be located in the same cluster (see SCode 1 and the description therein). The results however show that the average number of clusters with more than 1 active OR gene is consistent with the null model of random cluster allocation of OR genes (SFigure 10, p=0.112). Moreover, we found that the top two active OR clusters (ranked by expression level) always belonged to different chromosomes in each cell of the Late.INP stage. Thus the onset of OR gene expression in the multigenic stage cannot be caused by activation of an individual chromosome or a particular OR cluster. Conversely the transition to monogenic stage could be partially caused by restriction of expression to a single chromosome and cluster. However, this may not be the only selection mechanism involved, as some cells express up to 3-4 different OR genes simultaneously within a single cluster. Specifically, for the 34 Late.INP cells we found 20 cells, which expressed at least 2 OR genes from the same cluster (expression defined as ≥50 normalized counts). Thus, additional steps are required to restrict expression to a single gene within a cluster.

### Motif search in Greek island enhancers identifies novel transcription factors

Next we proceeded to identify potential factors involved in both onset of multiple OR gene expression and the transition to monogenic expression. We focused on transcription factors that had a detectable expression in our scRNA-seq dataset. We consider a factor detectable if it has at least 35 counts in at least 15 out of 294 cells in the neuronal lineage, leaving us with 1358 (co)TFs.

Previously, 63 intergenic enhancer regions, termed Greek islands, have been identified inside or near OR clusters using DNase I hypersensitivity-sequencing and chromatin immunoprecipitation sequencing^15^ and ATAC-seq^11^. SFigure 1 shows the co-localization of the Greek islands and the OR clusters on a map of the murine genome. Chromatin conformation capture experiments have revealed that Greek islands extensively contact OR clusters, remarkably both in cis and trans^38,39^.

We performed a de novo motif search on all Greek island enhancer regions as annotated by^11^ using MEME^40,41^ (Methods). Ungapped motif analysis of Greek islands identified one known motif, TYCCYWKGGGVCTHATTARM (reported in Monahan et al^11^), and two novel motifs GVDHCYYCAGRGAV and TBYTCHTCTCYMCAGDGWBNY, with E-value 1.7e^-057^, 3.7e^-027^ and 4.1e^-008^, respectively. Almost all Greek islands contain each of these motifs exactly once, except 8 Greek islands which are missing the third motif. TOMTOM was employed to align these motifs with known transcription factor motifs from the JASPAR database^42^ (Methods). TOMTOM did not predict any significant TF binding for the two novel motifs, therefore we do not discuss them further. We found 65 significant target binding sites for transcription factors inside the first motif (see Excel file 3). From those, 9 TFs were expressed at a detectable level.

The most significant motif, TYCCYWKGGGVCTHATTARM is composed of two adjacent submotifs, which are overlapped by a third submotif (Figure 3A). The first submotif is targeted by the COE1 DNA-binding domain which is found exclusively in the Ebf transcription factor family (Ebf1-4). The second submotif is bound by homeodomain TFs such as Lhx2, Emx2 and Uncx (Figure 3A). These results are consistent with previously reported Ebf1 and Lhx2 motifs to be positioned next to each other in most Greek islands^11^. Furthermore, the second submotif is expected to interact with transcription factors from three other families, the homeobox domain TF family, the Pou TF family (Pou6f1, this family has a strong enrichment in OR genes) and the ARID (AT-Rich Interaction Domain) domain TF family (Arid3a).

**Figure 3:**
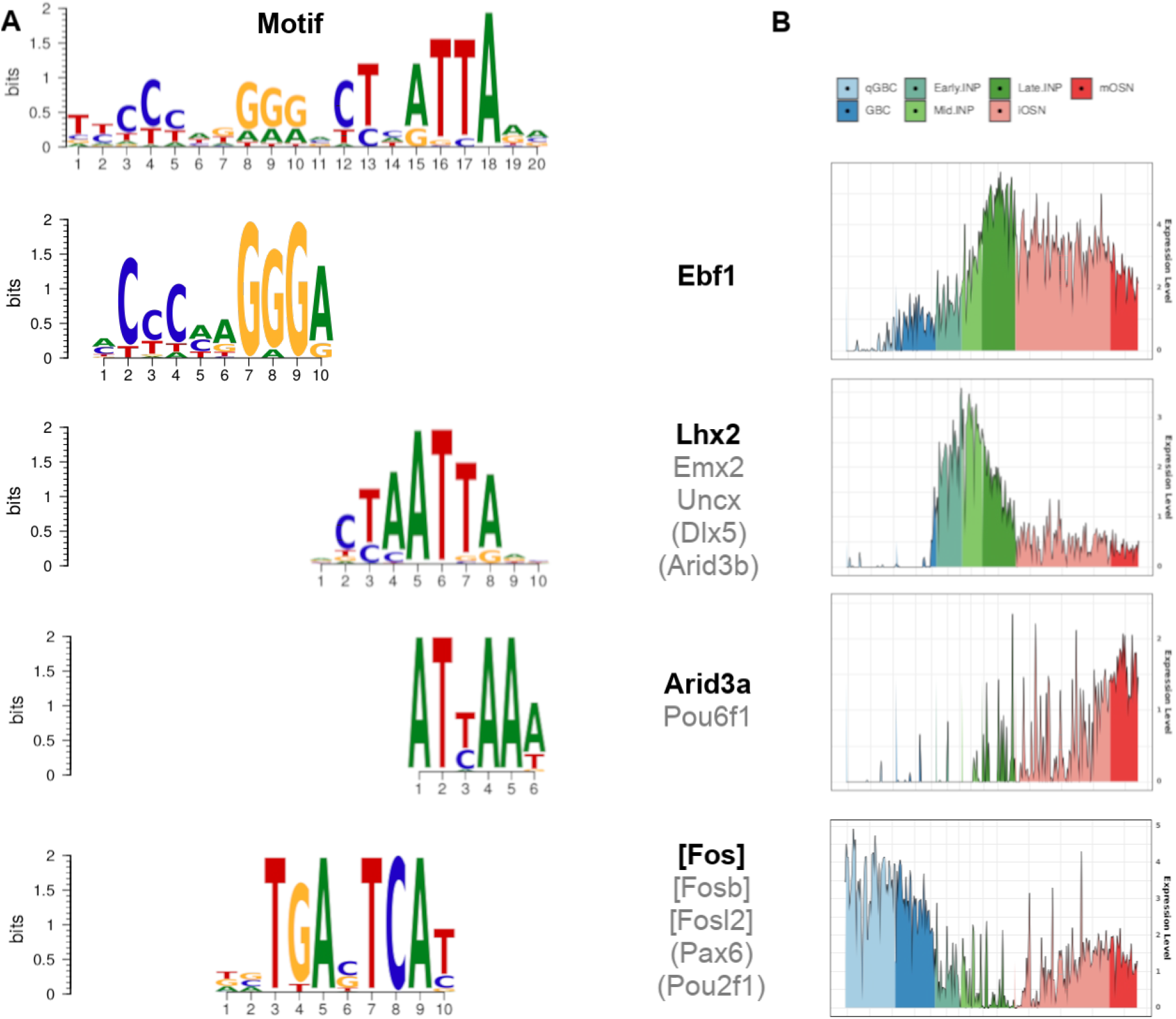
Greek islands binding motifs and representative pseudotime courses of respective TFs. **A)** Binding motifs found in Greek islands. The top row shows the motif found de novo. In the rows below, known binding sites of transcription factors that partly align with the de novo motif in different sites (TOMTOM) and/or are enriched in Greek islands (AME). The transcription factors given in square brackets refer to the TFs found only by Tomtom alignment whereas the round brackets refer to TFs found only by forward motif search (see SFigure 5). **B)** Single cell pseudotime courses of the TFs in bold print show characteristic trends along OSN differentiation. The grouping of the TFs in gray follows their pseudotime profile. It agrees with their grouping according to binding sites, except for Pax6 and Pou2f1 that have a homeobox domain like Lhx2.

Noteworthy, our analysis predicted a third submotif, which is a possible binding site for Fos, Fosb and Fosl2. Fos and Fosb are well-known early response transcription factors, which in turn regulate a broad variety of other transcription factors thus regulating many physiological processes^43–45^. This Fos-binding motif overlaps with the end of the first and the beginning of the second submotif, suggesting a cooperative/inhibitory interaction of the respective binding factors. This possibility will be investigated further in the context of the pseudotime analysis.

Complementary to the de novo motif analysis, we did a forward motif search with AME^46^ looking for known TF binding sites enriched in the 63 Greek Islands (Methods). This returned a total of 120 TFs (Excel file 4), of which 10 had a detectable expression (at least 35 count per cell in 15/294 cells of neuronal lineage) in our scRNA-seq dataset (SFigure 5). Six of those are also detected in the de novo motif search (see above), the additional TFs are Pax6, Dlx5, Pou2f1 and Arid3b (SFigure 3A). Dlx5 is part of the same TF family as Lhx2, which was found in the de novo motif search. Pax6 and Pou2f1 have a homeobox domain, whereas Arid3b is part of the same TF family as Arid3a.

Taken together we describe 13 TFs that are found at detectable levels and predicted to bind to Greek islands by de novo and/or forward motif search. Next we determined the pseudotime profiles of these TFs and found clear and distinct temporal expression patterns, providing additional evidence for their active involvement in the regulation of OR expression.

### Pseudotime analysis suggests transcription factors involved in OR expression

The sorting of cells according to pseudotime (Methods) generates, for each gene, a time course of its expression (see above). Notably, all TFs found by motif search in the previous paragraph show a pronounced temporal expression pattern, which belongs to one of three groups (Figure 3B and SFigure 5): The first group is active early in the silent phase, but strongly downregulated in late silent phase to reach a minimum in the multigenic phase (Fos, Fosb, Fosl2, Pax6 and Pou2f1). Some, but not all, are upregulated again in the monogenic phase (Fos, Fosb).The second group peaks within the multigenic phase (Ebf1, Lhx2, Emx2, Uncx and Dlx5). The third group is specifically upregulated during the monogenic phase (Arid3a and Pou6f1). Hereafter we will refer to these group definitions.

The fact that all TFs with a known Greek island binding site show a clear temporal pattern prompted us to perform a systematic search for TFs that change their expression upon transition between the three phases of OR expression. We also include co-factors in this analysis, because co-factors such as LDB1 have been found to be selectively associated with Greek islands and were suggested to initiate OR expression^38^. We searched for (co-)TFs with a significant differential gene expression of at least 2-fold between silent vs. multigenic phase or between multigenic vs. monogenic phase (Methods), resulting in 83 (34 going up, 49 down) respectively 39 (8 up, 31 down) relevant hits (Figure 4A, see Excel files 5,6).

**Figure 4.**
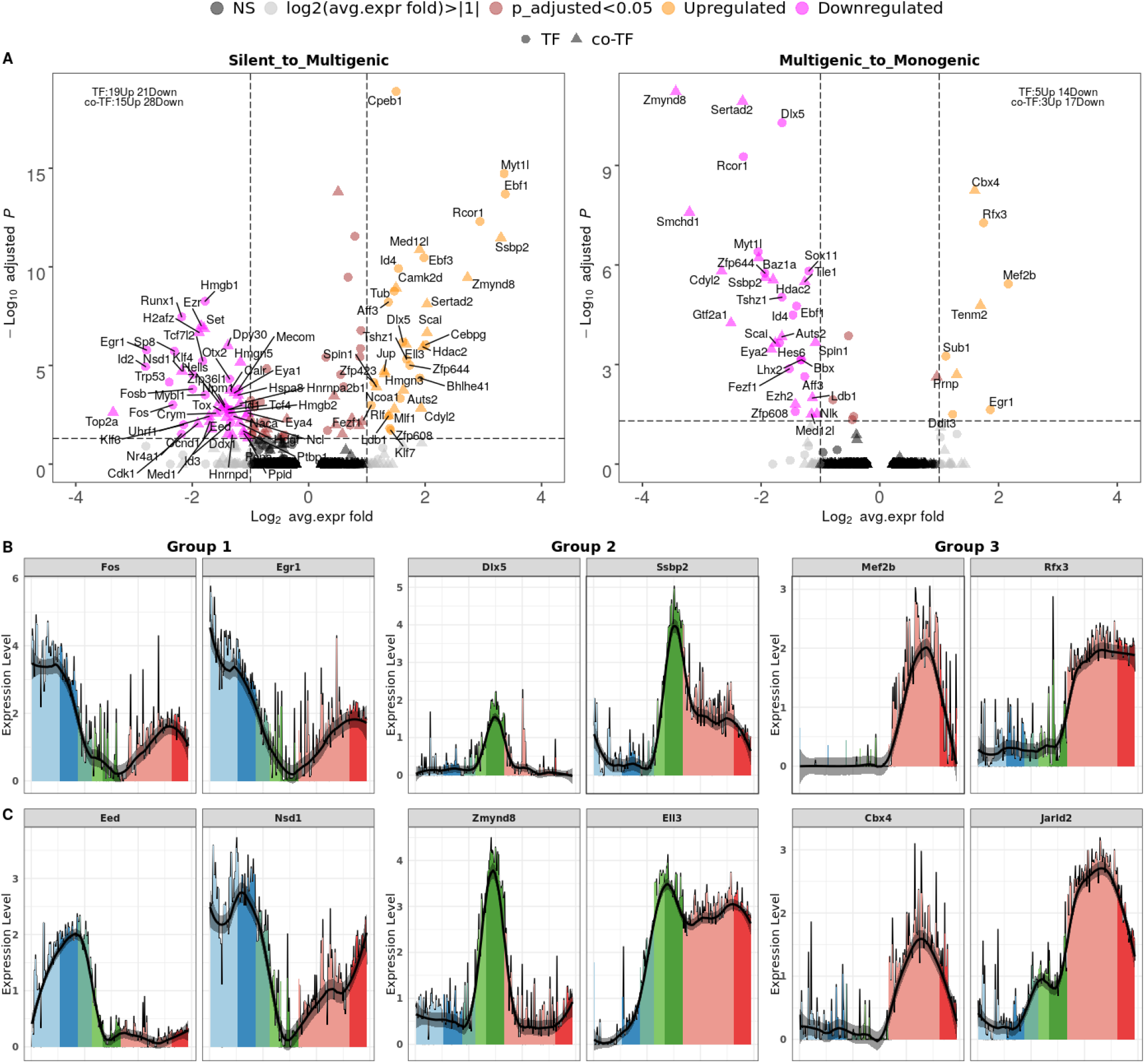
Differential expression analysis and pseudotime expression profiles of candidate factors. **A)** Volcano plots revealing the up- (peach) and downregulated (magenta) factors in the two transitions from silent to multigenic (left) and from multigenic to monogenic phase (right). TFs are shown as filled circles, cofactors are triangles. We selected (co-)TFs* with an adjusted p-value less than 0.05 (y-axis, log_10_ Bonferroni adjusted p-value) and an average expression change of at least 2 fold (x.axis, log_2_ fold). (Co-)TFs with only significant changes are shown in brown, those with only relevant expression fold > 2 are colored in grey, all others are in black (not significant). **B)** Single cell pseudotime courses of selected transcription factors and cofactors that show characteristic trends along OSN differentiation (cell stages are indicated by colors). Left to right: each group of 2 columns shows examples of transcription (co-)factors in group 1, 2, and 3 as defined in the text. **C)** Same as **B)** for chromatin remodelers. *Note that co-TFs in **A)** include chromatin remodelers.

Several of these differentially expressed factors were also identified by motif analysis (e.g., Ebf1, Lhx2, Dlx5, Fosb and Fos), but many are not. We manually inspected the pseudotime patterns of all differentially expressed (co-)TFs, and for detailed discussion, we selected those factors whose pseudotime expression pattern falls clearly into one of the three groups described above. We limit ourselves in the following to discussion of novel (co-)factors with additional evidence from motif analysis or a previous link to OR expression^11,18,19,38^. All other factors with a characteristic expression time course are shown in SFigure 9A.

For group 1 (active early in silent phase, downregulated in late silent phase and minimum in the multigenic phase) the differential expression search newly identified Egr1, whose expression resembles that of Fos and Fosb, which were already identified by motif analysis (see above). Therefore we searched explicitly for the known Egr1 binding motif (Methods) in Greek islands, which could be identified in 16 of 63 Greek islands. Fos, Fosb and Egr1 are immediate early genes, which are rapidly upregulated in response to external stimuli, immune response, and cellular stress^45^. Egr1, Fos and Fosb are specifically downregulated during the multigenic phase (Figure 4B and SFigure 9A). This suggests combinatorial interactions with the other components that regulate OR expression and will be discussed later.

The second group peaks specifically within the multigenic phase and 6 factors have been identified by motif analysis (see above). Differential expression analysis further obtains the cofactor Ssbp2 (Figure 4B) and three factors, Cebpg, Rcor1 and Ldb1, which have been reported previously to be involved in OR expression^11,38,47,48^. Ssbp2 binds to Ldb1 and thereby prevents Ldb1 from degradation^49,50^. While Ldb1 and Lhx2 were shown to bind to Greek island enhancers to regulate OR expression in trans^38^, Ssbp2 is a novel candidate with such a function.

The third group is specifically upregulated during the monogenic phase and two factors from this group have been identified by motif analysis. Differential analysis identifies additionally the TFs Mef2b, Rfx3 and Sub1, also known as PC4 (Figure 4B and SFigure 9A). Among these factors, only Rfx3 has a known motif, for which we performed a strict motif search in Greek islands (Methods). We report that 56 out of 63 Greek islands contain the binding motif for Rfx3. Moreover, note that Mef2a, which shares a similar SRF binding domain with Mef2b^51^, is found to be strongly bound to OR promoters^19^.

Taken together, our pseudotime analysis recovers a large proportion of candidate (co-)TFs identified by motif analysis - both for initiating onset of OR expression, and for the transition to monogenic stage. Moreover it extends the range of candidates whose time course correlates with these two transitions, and consequently the regulatory repertoire for these transitions.

### Changes in chromatin remodeler expression accompany both transitions in the OR selection process

It is known that chromatin changes accompany the selection of OR genes^20,21,23^. We therefore searched our data for chromatin remodelers that show expression changes during OR selection (Methods, Excel files 5,6). We confirmed previous observations that the chromatin remodelers Lbr and Cbx5 (SFigure 9B) are expressed at earlier stages and are downregulated in the course of OSN differentiation^15,21^. Furthermore, we discovered novel candidates for silencing OR genes, for onset of (multigenic) expression, and for transition to monogenic expression (Figure 4C):

Among the genes whose expression profiles fall into group 1 (minimum in multigenic phase), we found Eed, one of the constitutive subunits of the polycomb repressive complex 2, PRC2 (Figure 4C). Eed is required to maintain repressive H3K27me3 marks^52,53^ and its downregulation may lead to de-repression of OR expression in Late.INP stage. Note that another PRC2 subunit, Ezh2 is expressed during the silent phase as well, but decreases later, at the transition to monogenic phase (SFigure 6). Nsd1 is a histone methyltransferase that demethylates H3K36me2^54^ (Figure 4C). All remodelers found with a group 1 pseudotime profile (Hells, H2afz and Set) are predicted to play a repressive role in the silent phase of OR selection (we only show H2afz as example SFigure 9B).

For group 2 (peak in multigenic phase), we found prominent chromatin remodelers such as Zmynd8, Ell3, Sertad2, Med12l and Scai (Figure 4C, SFigure 9B). We also investigated the expression profile of Kdm1a which was known before as a regulator of OR expression. Kdm1a alias LSD1 is a Lysine demethylase and functions both as a coactivator by demethylation of mono-or di-methylated H3K9 and as a corepressor through demethylation of mono- or dimethylated H3K4^55–58^. There have been contradictory reports on the function of Kdm1a in OR expression as an activator^27^ or repressor of transcription^48^. The present data sheds light on this debate: While Kdm1a expression sharply peaks directly before the multigenic phase (arguing for its role as activator), it can be part of the Co-REST repressor complex^59^. Two components of the Co-REST repressor complex, Rcor1 and Hdac2, sharply peak during multigenic phase (Figure 5C, SFigure 8A), arguing for a change of function of Kdm1a by recruitment to the Co-REST complex at the transition to monogenic phase^55,60,61^.

**Figure 5.**
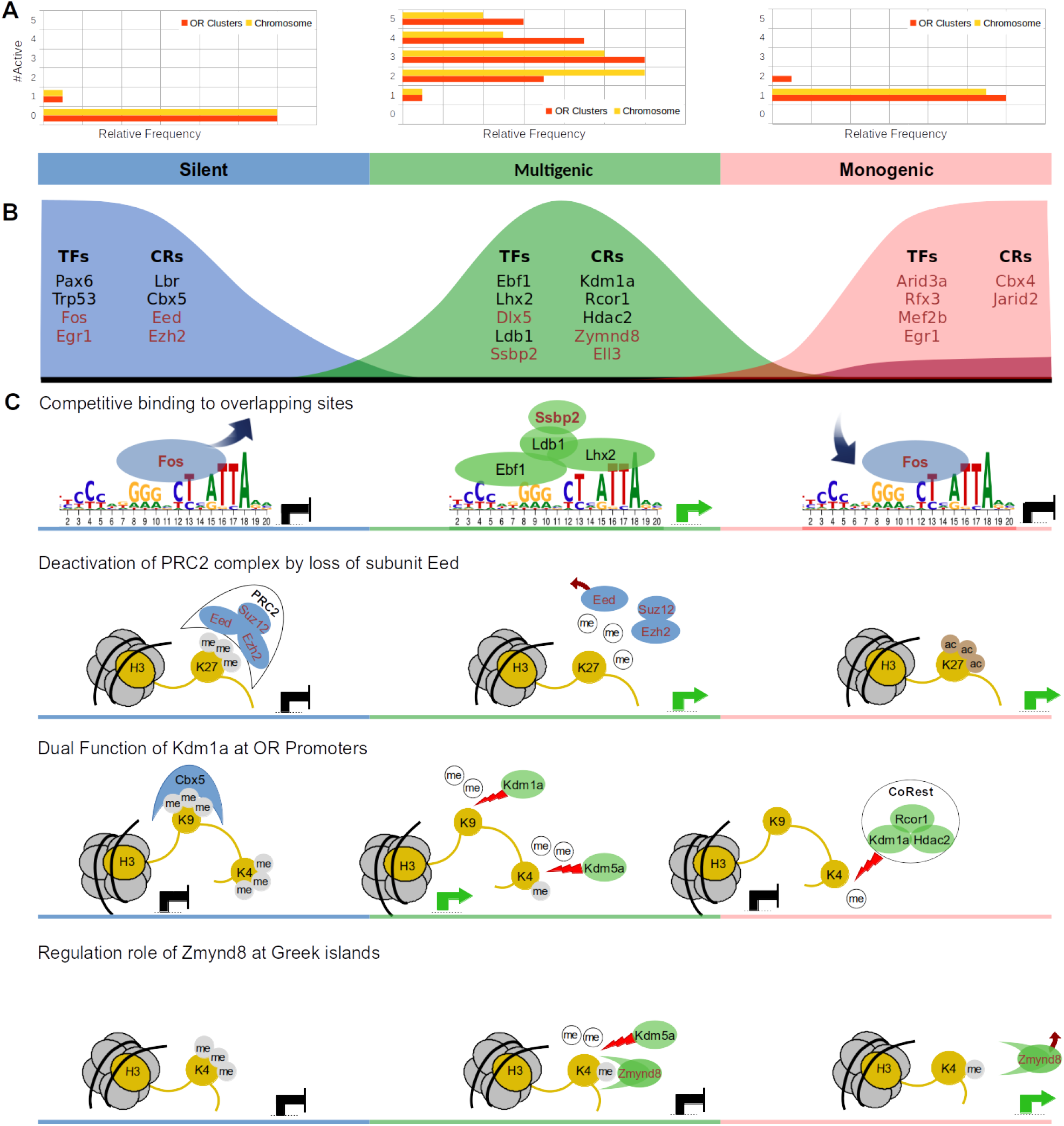
Graphical summary of OR gene expression and hypothetical selection mechanisms. **A)** Frequency distribution of active OR clusters and chromosomes per cell in the three phases of OR expression. Two exceptional cases of cells in multigenic phase with 6 respectively 7 active OR clusters are omitted from the plot. **B)** Pseudotime ordering of (co-)TFs and remodelers according to peak of expression in one of the three phases. Factors that so far have not been reported are highlighted in red. **C)** Four mechanisms potentially contributing to OR selection. Factors are colored according to their phase of peak expression. The placement of the nucleosomes is according to the pseudotime at which the respective processes take place. Green arrows indicate an activating effect on target genes, black blunt end arrows indicate an inhibitory effect. Top row: Competitive binding of Fos vs. Ebf1 and Lhx2 to different parts of the Greek island consensus motif, resulting in different enhancer activities during the three OR selection phases. Second row: Downregulation of the Eed subunit dissolves the PRC2 complex at the beginning of the multigenic phase and thus enables access of demethylases to H3K27me3. Third row: Methylated H3K9 is recognized, bound and protected by Cbx5 during the silent phase. At the end of this phase Cbx5 is downregulated and Kdm1a is upregulated. It participates in the process of H3K9 demethylation by removing the H3K9me1/2 mark. Later, after H3K4me3 has been demethylated to H3K4me2, presumably by Kdm5a during the multigenic phase, Kdm1a has a second role as part of the CoRest complex and demethylates H3K4me2. Bottom row: After demethylation of H3K4me3/2 by Kdm5a at Greek island enhancers, the chromatin reader Zmynd8, which peaks during the multigenic phase, can recognize H3K4me and repress enhancer activity. Activity is restored upon downregulation of Zmynd8 in the monogenic phase.

Of the four novel remodelers with a group 2 pseudotime profile, Zmynd8 and Ell3 are highly differentially expressed (Figure 4C). Zmynd8, a chromatin reader, is a particularly appealing candidate, since it is also known to play a role in the selective expression of the immunoglobulin heavy chain (*Igh*) regions in immune cells (B cells). Its product ZMYND8 binds *Igh* super-enhancers known as 3’ regulatory region (3’RR). ZMYND8 thereby controls the 3’RR activity by modulating the enhancer transcriptional status^62^. Consistent with an activating role during the multigenic phase, Ell3 does not only bind to active enhancers, but also marks the enhancers that are in a poised or inactive state in ES cells^63^.

Remodelers whose pseudotime profiles fall into group 3 (specifically upregulated during the monogenic phase) are *Cbx4* (Chromobox 4) and *Jarid2* (Figure 4C). Cbx4 is a component of a Polycomb group (PcG) multiprotein PRC1-like complex, which is required to maintain the transcriptionally repressive state of many genes^64,65^. *Jarid2* (Jumonji and AT-Rich Interaction Domain Containing 2) is required to repress expression of cyclin-D1 (CCND1) in cardiac cells by setting H3K9 methylation marks^66^, and it is upregulated upon transition from Late.INP to iOSN (SFigure 4B), i.e. upon exit from the cell cycle.

So far, we identified several chromatin remodelers that add to the regulatory repertoire for the onset of OR expression, and for the transition to monogenic stage. Another important feature which requires chromatin remodeling is the monoallelic expression of ORs in mature OSNs. This feature appears to be established from the very beginning of OR expression^4,23^. Factors involved in generating allelic exclusion therefore would be expected to peak at least as early as factors regulating the onset of (multigenic) expression (group 2 factors). We found two remodeling factors with a very early onset within group 2 which could potentially play such a role: *Smchd1* (structural maintenance of chromosomes flexible hinge domain-containing protein 1) and *Cdyl2* (chromodomain Y-like protein 2) (see SFigure 7).

Based on the interactions and temporal coordination of remodelers and (co-)TFs, we generate and discuss some hypotheses about OR selection below.

## Discussion

The monogenic expression of ORs presents a big challenge for the olfactory system, since it requires a random selection of exactly one OR per cell from the large family of OR genes. OR genes are known to be aggregated in silenced chromatin clusters during the silent phase, before the selection is made^21^. The identification of a multigenic state in early immature OSN^29,30^ by scRNA-seq made clear that the escape from silencing is not limited to a single OR gene in a single cell. In a single cell, we observed up to seven different OR clusters and more than a dozen different OR genes concomitantly active. Both transcription factors and chromatin remodelers have been identified as regulators of OR expression.

Here we have employed pseudotime analysis of a single cell transcriptome data set^31^ focusing on OR expression. We aligned the three main phases of OR expression, silent, multigenic and monogenic^29,30^, with the stages of OSN differentiation as defined in^31^. By analysis of winner vs. runners-up OR expression, we could precisely assign the onset of multigenic phase to Late.INP, and the onset of monogenic selection to iOSN. Most cells in the multigenic phase express ORs from more than one cluster and more than one chromosome (Figures 2B, 5A). We tested whether ORs co-expressed in one cell tend to lie in identical OR clusters and did not find evidence for that (SCode 1, SFigure 10). We conclude that the selective expression of OR genes during multigenic phase is not caused by escape from compaction of merely one OR cluster. In the monogenic phase, the vast majority of cells express only a single OR gene (Figure 5A).

We then identified candidate factors (TFs and cofactors including chromatin remodelers) differentially expressed between these stages and thus potentially involved in the transitions silent-to-multigenic (40 TFs, 43 cofactors) and multigenic-to-monogenic (19 TFs, 20 cofactors) (Methods, Excel files 5,6). Many of these differentially expressed factors are likely not directly involved in OR selection, since cells undergo many substantial changes along OSN differentiation. Thus, we performed an independent de novo motif analysis of Greek islands, which should enrich TFs involved in OR selection. This search revealed one motif that could be decomposed into three consecutive submotifs, only two of which were described previously^11^. We additionally recognized a central Fos binding motif overlapping the previously described two elements (Figure 3A). All but one of the factors that bind to these motifs show characteristic pseudotime expression time courses, which could be classified in three groups (Figure 5B). The temporal coordination of these factors together with the precise location of their respective binding sites enables us to generate hypotheses about their molecular interactions and possible functional consequences.

In the silent phase the transcriptional regulator Fos is binding to the central motif and may competitively prevent binding of the known activators of OR expression, Lhx2 and Ebf1^11^, which bind to the left and right submotifs (Figure 3A, top row of Figure 5C). The strong downregulation of Fos expression in the multigenic phase then would allow Lhx2 and Ebf1 to access their binding motifs, and recruit their known binding partner Ldb1 to Greek islands^38^. Ldb1 mediates trans interactions between different Greek islands, creating super-enhancer hubs that include neighboring OR clusters^38^. Our pseudotime analysis predicted additionally the co-factor Ssbp2 to play a role in this process. Ssbp2 is in other contexts known to bind to Ldb1 and thereby prevents its degradation by the proteasome^49,50^. Thus expression of Ssbp2 in the multigenic phase would amplify the effect of Ldb1.

All four factors peak during the multigenic phase (Figure 4A-B, Figure 5B and SFigure. 9A). This expression is anti-correlated to that of Fos, which supports the competitive interaction hypothesis outlined above (first row of Figure 5C). The high activity of Fos in silent and monogenic phase would counteract complex formation even of the small concentrations of Lhx2 and Ebf1 observed in this stage.

Our results found no OR expression during the Mid.INP stage despite significant expression of Lhx2, Ebf1 and Kdm1a (known activators of OR genes). We note that all components of the polycomb repressive complex 2 (PRC2), Eed, Ezh2 and Suz12, are active in Mid.INP, which could explain the absence of OR gene expression in Mid.INP despite the presence of the activators (second row of Figure 5C). Moreover, an essential subunit of PRC2, Eed, is significantly reduced during onset of OR expression in Late.INP (Figure 4 and SFigure 6). This elimination of Eed is sufficient to disassemble the PRC2 complex, which then can no longer maintain the repressive H3K27me3 mark^52,53^. Furthermore, we showed a dramatic reduction in expression of Ezh2 and Suz12 subunits of PRC2 along OSN differentiation (second row of Figure 5C and SFigure 6). We conclude that PRC2 activity may be involved in repression of OR expression in Mid.INP. The disassembly of PRC2 in Late.INP then enables the Greek Island hubs to transiently activate the cis-corresponding OR gene/s, which enables the expression of multiple OR genes in most of Late.INP stage cells at the same time with relatively low levels compared to later stages of OSN differentiation (monogenic phase).

Heterochromatic silencing of ORs throughout OSN differentiation is enforced by the (interchromosomal) convergence of OR loci to OSN-specific, highly compacted nuclear bodies^21^. It has been shown that in the monogenic phase individual active OR genes require de-silencing by lysine demethylase Kdm1a^27^ and spatial segregation of the single chosen OR allele towards euchromatic nuclear territories^21^. Another study however showed that the deletion of Kdm1a leads to persistent multigenic expression, suggesting a silencing role of Kdm1a rather than activating one^48^. Here, pseudotime analysis sheds light on the seemingly contradictory role of Kdm1a (third row of Figure 5C):

The CBX5 protein is responsible for silencing of OR genes during the silent phase by binding to and thereby protecting the repressive H3K9me3 mark in gene bodies^21^. It vanishes upon transition to multigenic phase (SFigure 9B). After H3K9me3 has lost one methyl group (e.g., through the action of Kdm4a), Kdm1a can demethylate H3K9me2. Kdm1a peaks at Mid.INP stage and acts on di-methylated lysines only^55^. Thus the action of Kdm1a on H3K9me3 leads to activation in Mid.INP (Figure 5C).

In contrast, another methylation site, on H3K4 is a mark of an active promoter/enhancer in the *methylated* stage (trimethylated for promoter, monomethylated for enhancer). Kdm5a peaks during multigenic phase (Figure 5C, SFigure 8A) and can demethylate tri- or di-methylated H3K4 to its monomethylated form. Kdm1a by itself cannot act on H3K4me, but in a complex with Rcor1 and Hdac2 (CoRest complex) it is able to demethylate H3K4me2/1^55,60,61^(Figure 5C). Rcor1 and Hdac2 have their peak expression during Late.INP stage, i.e. shortly after onset of the Kdm1a peak (Figure 5B, SFigure 8A). Thus, in Late.INP but not in Mid.INP, Kdm1a can demethylate H3K4me2/1, resulting in repression. This amounts to the multigenic phase (Late.INP) beginning to build up the molecular machinery to downregulate all but one of expressed ORs.

Taken together, the same enzymatic activity of the same factor (demethylation by Kdm1a) results in opposing effects on transcription due to co-factors Rcor1 and Hdac2 modulating substrate specificity of Kdm1a. Moreover our pseudotime analysis supports the hypothesis that OR expression issues a negative feedback signal on Kdm1a which is mediated by Atf5 and Adcy3 during transition from multigenic to monogenic phase^25,27,28^ (see SFigure 8B).

During the monogenic phase, a super-enhancer is formed by the trans interaction between multiple Greek islands. Among the OR genes associated with this super-enhancer, merely one OR is expressed at very high levels^38^. During the multigenic phase, H3K4me3 marks of Greek islands^20^ are converted to H3K4me, e.g. by Kdm5a (SFigure 5C). The latter histone mark can be recognized by the chromatin reader Zmynd8^67^. Zmynd8 has been shown to participate in another, highly specific selection process, namely the expression of *Igh* genes^62^. There, it recognizes H3K4me and represses super-enhancer activity in B cells. We therefore speculate that it might play a similar role in OR expression.

Expression of Zmynd8 peaks simultaneously with expression of the super-enhancer forming complex Ebf1-Lhx2-Ldb1-Ssbp2 during multigenic phase (Figure 4C for Zmynd8 and Ssbp2, Figure 3 for Ebf1 and Lhx2, SFigure 9 for Ldb1). Upon transition to the monogenic phase, Zmynd8 expression ceases while Ebf1 and Sspb2 maintain intermediate expression. The disappearance of the ZMYND8 protein might abolish the suppression of the super-enhancer and could allow very high expression of the selected OR (Figure 5C bottom row).

So far it is unknown whether the monoallelic expression characteristic for mature OSNs is preceded by a biallelic phase. The dataset evaluated here does not allow us to analyze this question directly, because it does not exhibit enough sequence diversity between alleles to distinguish them. We searched for epigenetic factors known to be involved in allelic selection in other contexts, which show significant and relevant expression changes during OR differentiation. We found two factors, Smchd1 and Cdyl2 (SFigure 6), which were discussed as stabilizing monoallelic expression. They play a role in epigenetic silencing, spermatogenesis, random inactivation of X chromosome, and stabilize monoallelic expression^68-70^. Both factors peak sharply in mid to late INP. Assuming these two factors initiate monoallelic expression, this would place monoallelic selection before or concomitant with the onset of (multigenic) expression in Late.INP, in other words, there might be no biallelic stage at all.

The dissection of the OSN maturation process into different stages allowed us to reveal three phases of OR gene selection. This in turn enabled an in-depth analysis of pseudotime expression profiles^71^, leading to several promising candidates and to testable hypotheses on the mechanisms involved in OR gene selection. It will be interesting to complement the present data with single cell chromatin accessibility data (e.g. scATAC-seq) or single cell chromatin conformation data (Hi-C) in the context of conditional knockouts of the factors we have identified. Experiments should be carried out in hybrid crosses to additionally monitor allelic expression. Our study has narrowed down the cell stages and the time window that need to be analyzed for these purposes, thereby enhancing future research on this topic.

## Methods

Most statistical analysis and visualization were done in RStudio using R version 3.6.3.

### a. Data processing and Quality control

Our analysis of OR expression patterns during OSN differentiation is based on a scRNA-seq dataset generated by (Fletcher et al^31^, GEO: GSE95601). Ngai group investigated the homeostatic differentiation in the postnatal olfactory epithelium. Horizontal basal stem cells (HBC) were released from quiescence by a conditional knockout of the Trp63 transcription factor. Briefly, cells were FACS (fluorescence-activated cell sorting) selected for Sox2-EGFP-pos-itive, ICAM1-negative, SCARB1/F3-negative expression to enrich for the cell population of interest (GBCs, later neuronal intermediates, and microvillous cells over sustentacular cells). Then scRNA-seq was done using the Fluidigm C1 microfluidics cell capture platform followed by Illumina multiplex sequencing. Fletcher et al^31^ used RefSeq transcript annotations to align reads to the GRCm38.3 mouse genome with Tophat2. This resulted in 849 cells with a mean coverage of 1.36M reads per cell (before read quality control). Contaminations, doublets and other technical artifacts were removed according to the ‘oeHBCdiff_filtering.R’, protocol (https://github.com/rufletch/p63-HBC-diff).

From here on our data processing differs from Fletcher et al^31^. We included all transcripts (above 40 counts in total) that occured in at least one cell to ensure retrieval even of OR genes with very sporadic expression. Further filtration was done using Seurat (version 3.1.4)^72–74^, keeping only cells with at least 1250 expressed genes. A total of 687 cells with a mean library size of 460k unique reads passed all filtration criteria and the median number of genes per cell is 4164. The library size distribution of these cells is shown in (SFigures 2,3).

Duplicate removal and normalizing the counts was performed by Seurat’s SCTransform with default parameters^72^, resulting in a 687 cells times 17228 genes matrix followed by Seurat workflow^72^.

### b. Dimension reduction, clustering and cell type assignment

We followed the Seurat clustering workflow. First, dimension reduction was done using Principal Component Analysis (PCA). The number of principal components kept was set to 15, after assessing the goodness of approximation by JackStraw and ElbowPlot functions. A shared k-nearest neighbor graph was built by the FindNeighbors function. Afterwards, the Lovain algorithm was applied to define 13 distinct clusters from the shared nearest neighbor graph using the FindClusters function and the Jaccard index as a similarity measure. This number matches the number of clusters identified in (Fletcher et al^31^) for this data. The expression of cell type marker genes that were collected from the literature (STable 1) served to assign cell clusters manually to known cell types according to the Seurat guidelines^72^. At least two expressed markers were required to confidently annotate a specific cell type. Visualization of the data was performed by PCA, UMAP^75^ and tSNE^76^.

### c. Trajectory inference and pseudotime assignment

Slingshot^36^ was used to construct a minimum spanning tree (MST) based on the 15-dimensional representation of the cells obtained above. The topology of the MST is independent of the root choice. For biological reasons, we selected qGBC as the root for the assignment of pseudo time, where it can differentiate to any cell type of MOE^35^. For each cell, a pseudotime between 0 (cells at the root node) and 1 (leaf node cells) was assigned by the slingPseudotime function.

### d. Differential expression analysis

For each cell stage, we identified marker genes showing differential expression compared to all other cell stages using FindAllMarkers in Seurat, using the Wilcoxon rank sum test. Supplemental Figure 3 shows a heatmap of the top 10 differentially expressed genes (i.e., putative markers) for each cell stage of MOE.

Among 2712 (co-)TFs (include chromatin remodelers) obtained from the GO.db package and AnimalTFDB3.0 (http://bioinfo.life.hust.edu.cn/AnimalTFDB/#!/), we found 2004 (co-)TFs expressed (at least one count in one cell) in the neuronal lineage. In later differential expression analysis we compared the expression profiles of cell stages that were placed consecutively along the neuronal trajectory (i.e., the maturation of OSN) to identify genes that change their expression upon transition between cell stages (SFigure 4B). Volcano plots for each cell type transition were generated by a slightly modified EnhancedVolcano function (https://github.com/kevinblighe/EnhancedVolcano).

Differential expression analysis for silent to multigenic and multigenic to monogenic phase transitions were performed by comparison of pre-Late.INP cells vs. Late.INP cells and Late.INP vs. post-Late.INP cells, respectively. Genes with a Bonferroni adjusted p-value <0.05 (Wilcoxon rank sum test, FindMarkers function in Seurat) and an average absolute FC >=2 were considered differentially expressed. This yielded 83 respectively 39 differentially expressed (co-)TFs for the two transitions.

### e. Motif Analysis

The genomic ranges of 68 OR clusters and 63 Greek islands were compiled from Monahan et al^11^, which allows the matching of OR clusters and Greek islands. UCSC genome browser tools were used to find all genes inside OR clusters. We performed a motif search on the 63 Greek island sequences (Fasta file 1) and the approximate promoter regions (500bp upstream the transcription starting sites) of all OR genes were obtained by using the “ucsc-twobittofa” bioconda package and the “biomart” R package respectively. The MEME suite web server for motif search and analysis^40,41^ was used to predict the transcription factors (TF) that bind to Greek islands. We applied MEME using default values for all parameters to find the novel, ungapped motifs inside Greek islands with the following command: meme greek_islands.fa -dna -oc. -nostatus -time 14400 -mod zoops -nmotifs 3 -minw 6 -maxw 50 -objfun classic -revcomp -markov_order 0

Then we performed motif comparison between each motif found in the above-mentioned analysis against a database of known TFs motifs (JASPAR2018_CORE_non-redundant and uniprobe_mouse databases) using Tomtom tool^42^. The Pearson correlation coefficient was used to measure the similarity between columns of position weight matrices (PWMs) and we restricted the results by setting q-value <= 0.1 (rather than 0.5 by default) as a threshold (10% FDR) using the following command: tomtom -no-ssc -oc. -verbosity 1 -min-overlap 5 -mi 1 -dist pearson - thresh 0.1 -time 300 query_motifs db/MOUSE/uniprobe_mouse.meme db/JASPAR/JASPAR2018_CORE_non-redundant.meme

We also investigated the enrichment motifs in 63 Greek islands sequences using AME tool^46^ by using an average odds score method and fisher’s exact test as a motif enrichment test through the following command: ame --verbose 1 --oc. --scoring avg --method fisher --hit-lo-fraction 0.25 --evalue-report-threshold 1.0 --control --shuffle-- --kmer 2 greek_islands.fa db/MOUSE/uniprobe_mouse.meme db/JASPAR/JASPAR2018_CORE_non-redundant.meme

Finally, a strict motif search in Greek islands for selected TFs was done by “ucsc-findmotif” bioconda package, allowing for 3 mismatches.

### f. Visualization of time series

Grouped time series^77^ was used to visualize pseudotime series of individual genes and to calculate and visualize aggregated groups of genes, e.g. all OR genes. Since the original expression count matrix is sparse (75.45% zero count entries), we first applied ALRA^78^, which has specifically been designed for the imputation of missing values in scRNA-Seq data (SFigure 3). The imputed expression matrix retrieved ~2403 missing values, reducing the fraction of zero count entries to 61.50%. The median number of expressed genes per cell was 6715 (see SFigure 2,3). Note that the imputed expression matrix was used only for visualization, for all analysis steps we used normalized counts without data imputation.

## Author contribution statement

A.T. and S.I.K. conceived and designed the analysis; M.H. collected the data, conducted the analysis and prepared the figures. All authors wrote, read and approved the manuscript.

## Data availability statement

The scRNA-seq dataset was obtained from: https://www.ncbi.nlm.nih.gov/geo/query/acc.cgi?acc=GSE95601

## Additional information

The authors declare no competing interests.

## Notes

https://www.ncbi.nlm.nih.gov/geo/query/acc.cgi?acc=GSE95601

### Competing Interest Statement

The authors have declared no competing interest.

