## Supplementary material for "Pseudotime analysis reveals novel regulatory factors for multigenic onset and monogenic transition of odorant receptor expression": Supplementary_Figures.pdf

### Abstract

During their maturation from horizontal basal stem cells, olfactory sensory neurons (OSNs) are known to select exactly one out of hundreds of olfactory receptors (ORs) and express it on their surface, a process called monogenic selection. Monogenic expression is preceded by a multigenic phase during which several OR genes are expressed in a single OSN. Here, we perform pseudotime analysis of a single cell RNA-Seq dataset of murine olfactory epithelium to precisely align the multigenic and monogenic expression phases with the cell types occurring during OSN differentiation. In combination with motif analysis of OR gene cluster-associated enhancer regions, we identify known and novel transcription (co-)factors (Ebf1, Lhx2, Ldb1, Fos and Sspp2) and chromatin remodelers (Kdm1a, Eed and Zmynd8) associated with OR expression. The inferred temporal order of their activity suggests novel mechanisms contributing to multigenic OR expression and monogenic selection.

### Supplementary Figures

#### Genomic locations of OR clusters and Greek islands

**SFigure 1:** Genomic locations of OR gene clusters (black bars) and Greek island enhancer regions (red bar) in each murine chromosome. Chromosomes are scaled to identical width for visualization purposes.

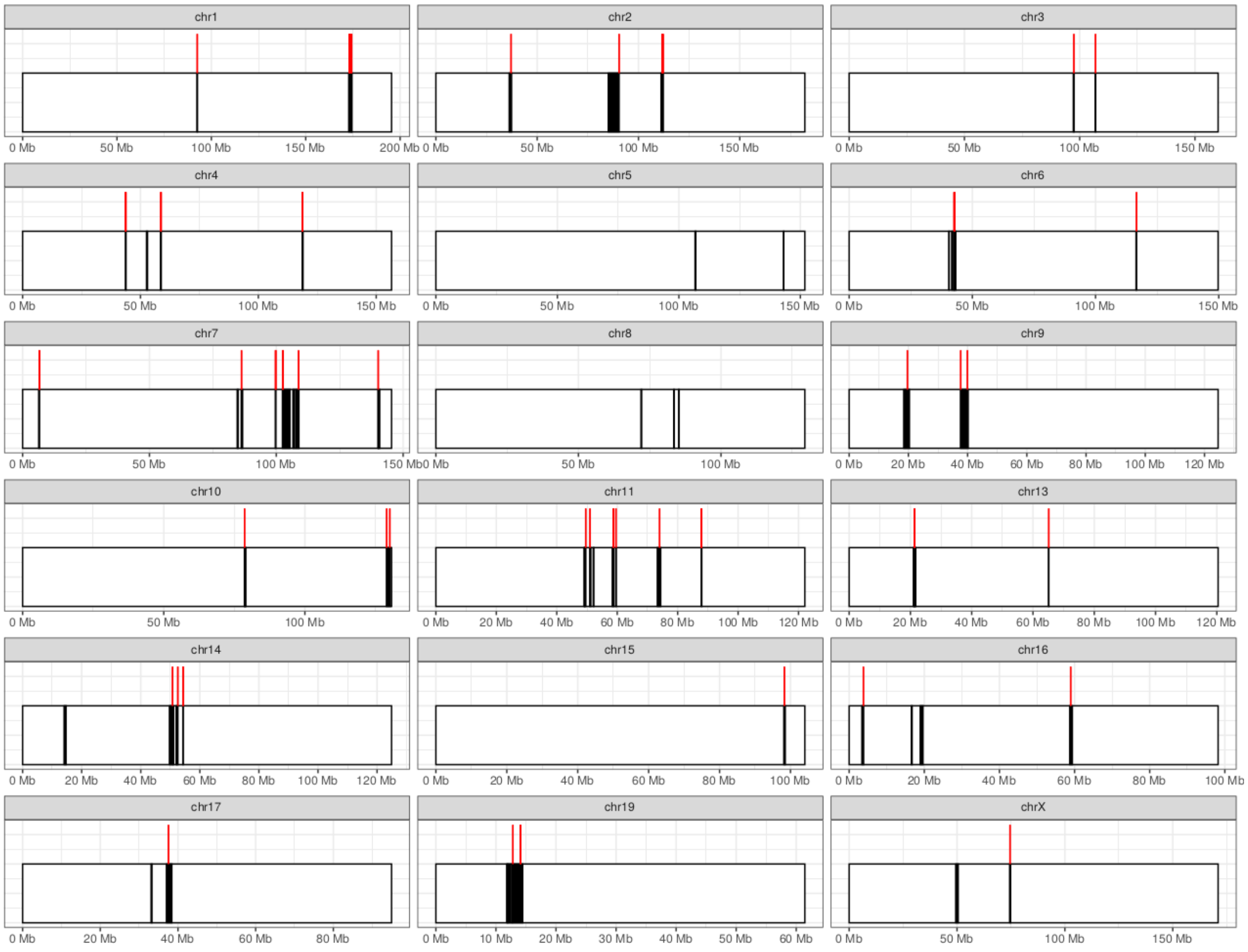

### Distribution of detected genes per cell before and after imputation

**SFigure 2: Library size per cell after final filtration and without imputation.**
**a)** Violin plot of number of expressed genes per cell in each cell type. **b)** Violin plot of number of reads (nCounts) per cell in each cell type where each black dot represents the library size of a single cell.

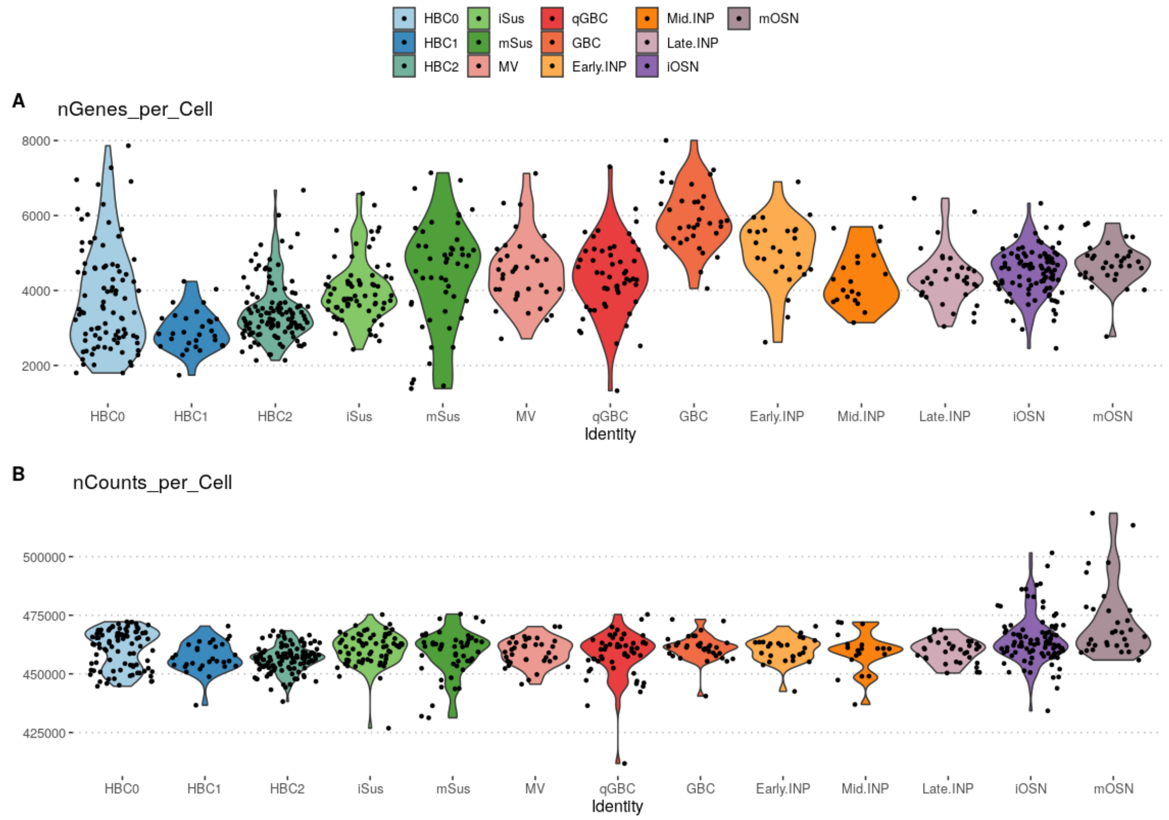


**SFigure 3: Distribution of genes per cell after imputation (dealing with dropouts).**

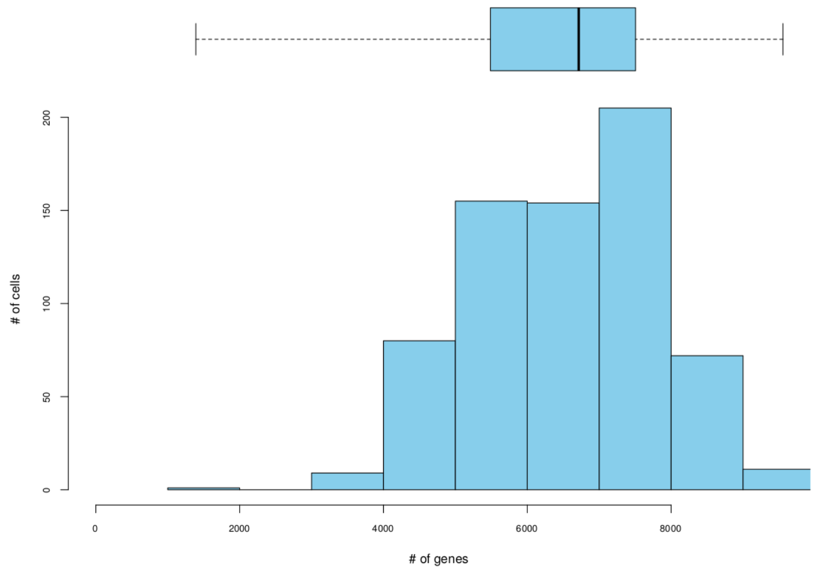

#### Cell type marker genes

**STable 1:** List of marker genes that we used to assign individual cells to cell types, according to their scRNA-Seq expression profile.

| **Cell type** | **Marker gene(s)** | **References** |
|---|---|---|
| Horizontal basal stem cells (HBC0, HBC1 and HBC2) | *Trp63, Krt5* and *Krt14* | Fletcher et al. 2017; Fletcher et al. 2011 |
| Sustentacular cells<br><br>(iSus and mSus) | *Cyp2g1, Notch2, Hey1, Sox2* and *Cyp1a2* | Rodriguez et al. 2008; Fletcher et al. 2017; Durante et al. 2020 |
| Microvilious cells (MV) | *Ascl3, Cftr* and *Coch* | Fletcher et al. 2017; Genovese & Tizzano 2018; Durante et al. 2020 |
| Globosal basal stem cells (qGBC and GBC) | *Lgr5, Tmprss4, Kit Mki67 and Top2a, Hes6* and *Ascl1* | Leung et al. 2018; Goss et al. 2016; Packard et al. 2016; Goldstein et al. 2015; Jang et al. 2014; Chen et al. 2014; Huard et al. 1998; Caggiano et al. 1994 |
| Immediate neuronal precursor 1 and 2<br><br>(INP.Early and INP.Mid) | *Neurod1, Neurog, Top2a, Mki67 and Lhx2* | Fletcher et al. 2017; Packard et al. 2016 |
| Immediate neuronal precursor 3<br><br>(INP.Late) | *Lhx2, Ebf1 and Gap43* | Fletcher et al. 2017; Packard et al. 2016 |
| Immature olfactory sensory neurons (iOSN) | *Gng8, Trib3 and Gap43* | Hanchate et al. 2015;Lin et al. 2017; Fletcher et al. 2017; Durante et al. 2020 |
| Mature olfactory sensory neurons (mOSN) | *Gng13, Gnal and OMP* | Hanchate et al. 2015;Lin et al. 2017;Fletcher et al. 2017;Durante et al. 2020 |

### DE Plots

**SFigure 4A:** Heatmap plot shows the expression profiles of top 10 differentially expressed genes in each cell type starting from HBC0 (top left) and ending with mOSN (bottom right). The gradient scale colors represent the normalized expression in log scale: white (no expression), blue (low to high expression) and red color (very high expression).

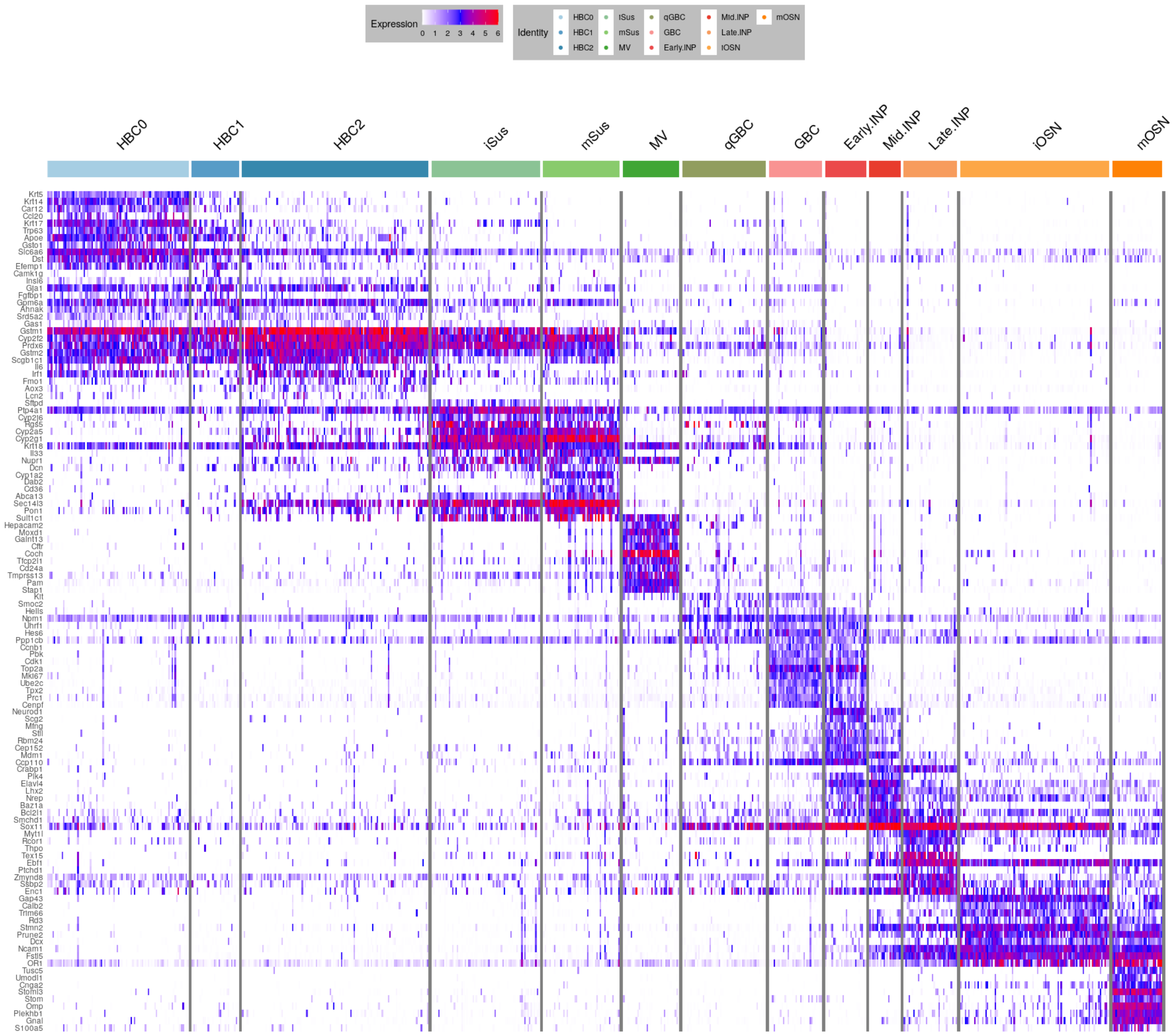

**SFigure 4B: The upregulated and downregulated (co-)TF genes from stage to stage along neuronal lineage.** We selected (co-)TFs with an adjusted p-value less than 0.05 (y-axis, $\log_{10}$ Bonferroni adjusted p-value) and an average expression change of at least 2 fold (x.axis, $\log_2$ fold).

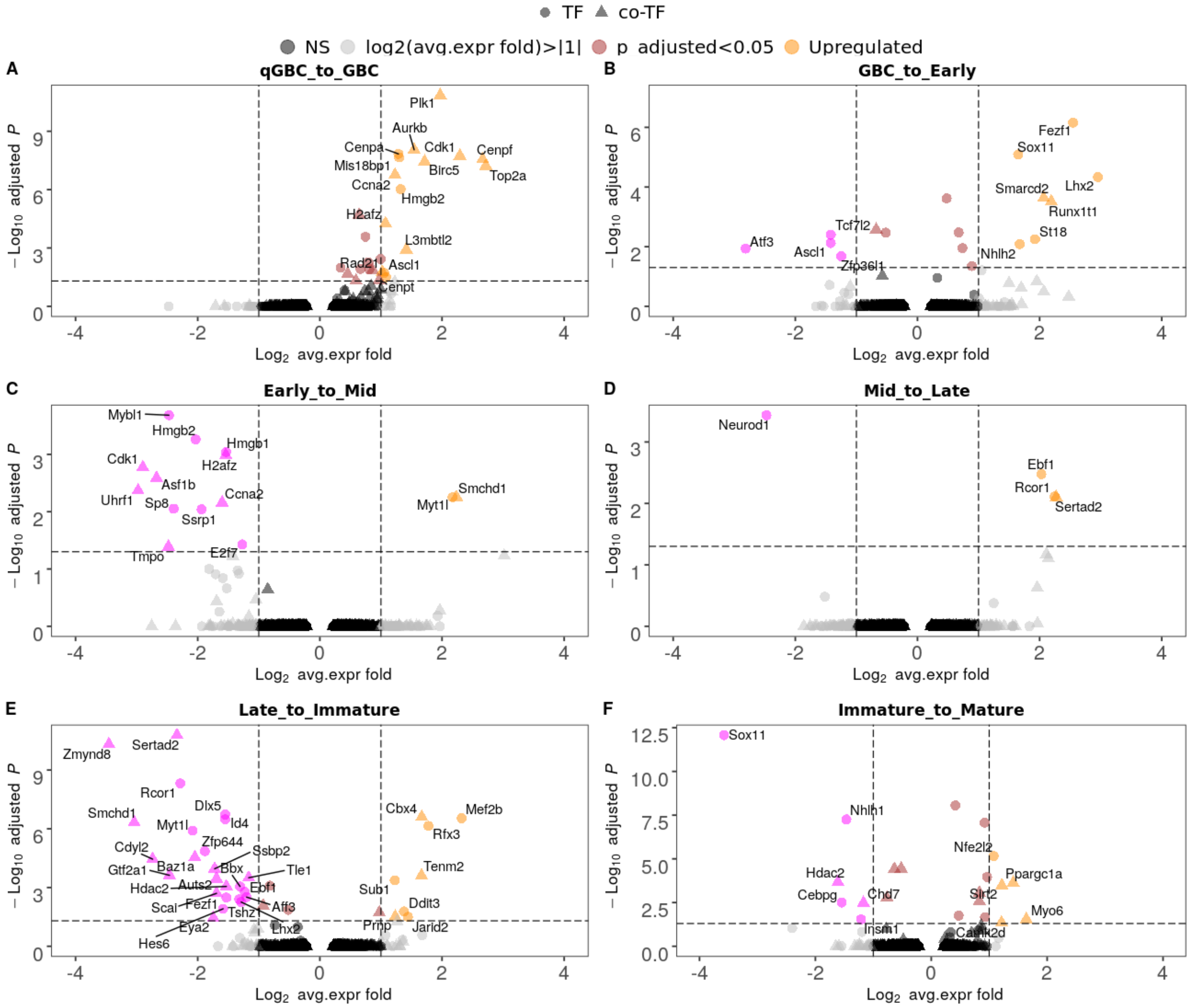

### VlnPlot of all TFs found in motif search

**SFigure 5**: (co)-TFs binding into Greek islands (Motif search results) that express at least 35 counts in at least 15 out of a total of 294 cells.

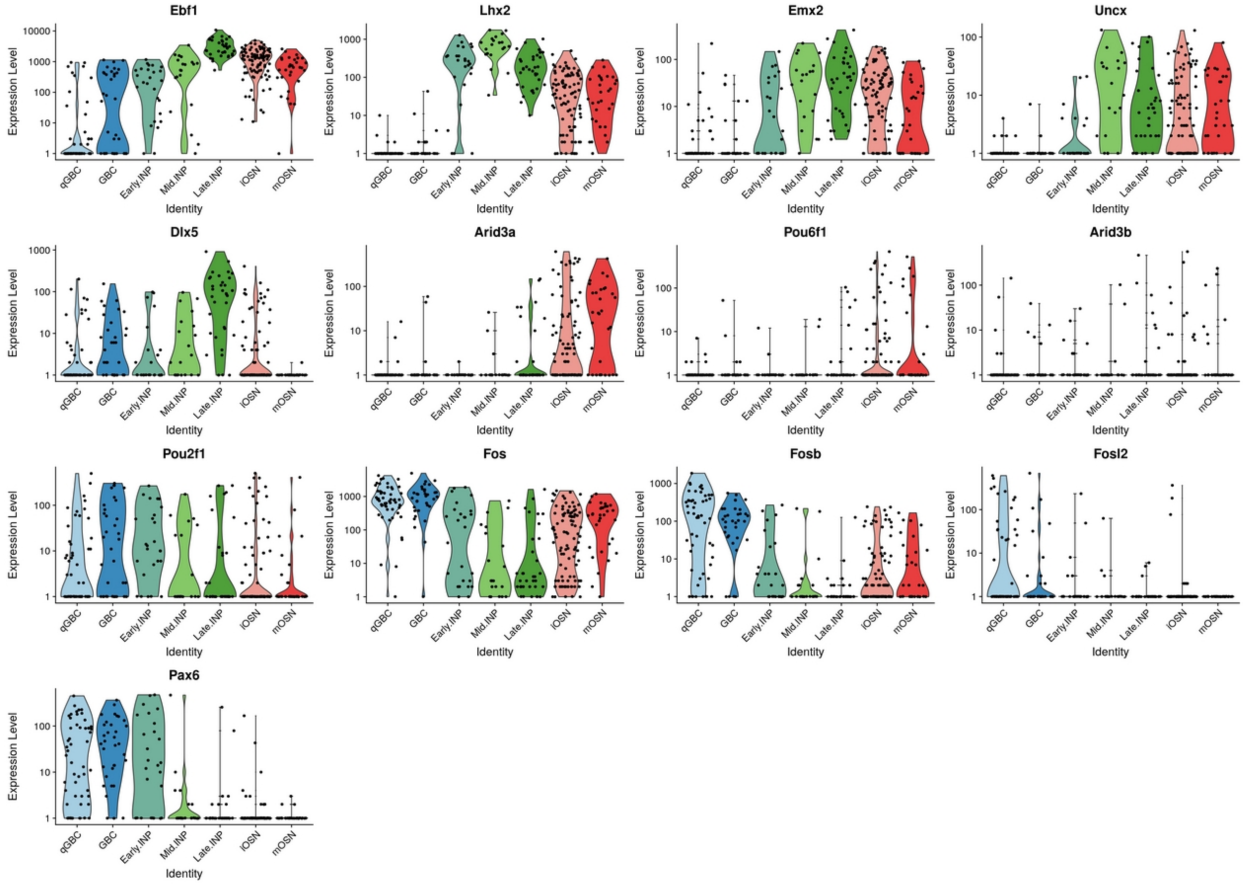

### PRC2 complex (H3K27 methylation)

**SFigure 6**: **Transcriptional dynamics of PRC2 complex.** The top row shows the pseudotime expression timecourse for each of the PRC2 subunits (Eed, Ezh2, Suz12) separately. The bottom row shows the expression distribution of each subunit per cell type. Numbers indicate the p-value (Wilcoxon rank sum test) of cell type transitions that show a significant expression change. Note that the expression of the Eed subunit is almost zero in Late.INP stage, indicating that the elimination of PRC2 is done during transition from Mid to Late.INP stage.

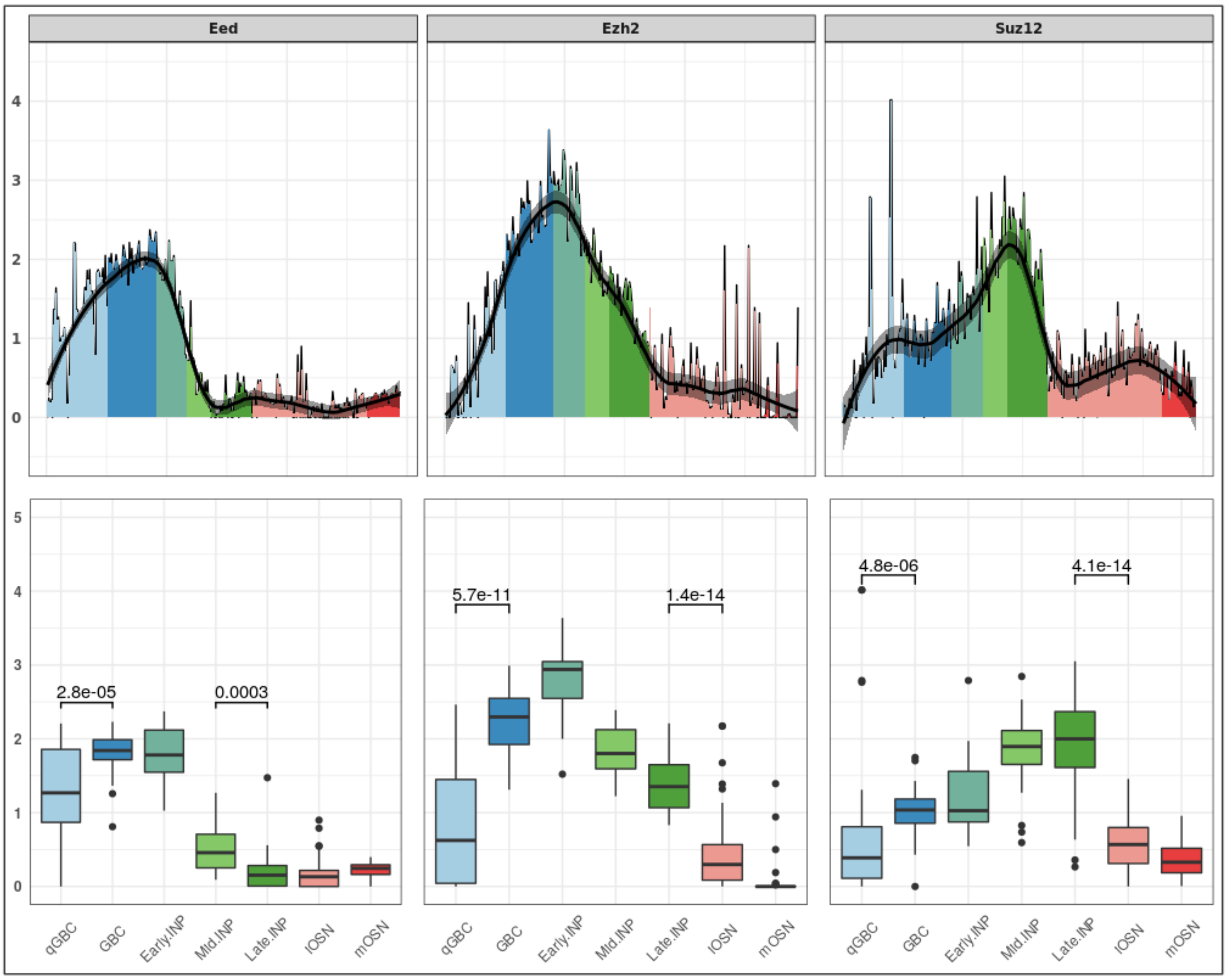

### Monoallelic factors

**SFigure 7: Monoallelic silencing factors.** The transcriptional dynamics of Smchd1 (light blue) and Cdyl2 (blue) show the same expression pattern along neuronal lineage, indicating that they probably are recruited for the same function (stabilizing the monoallelic selection after asynchronous replication of OR genes) during their peak period of neuronal lineage (Mid and Late.INP stages).

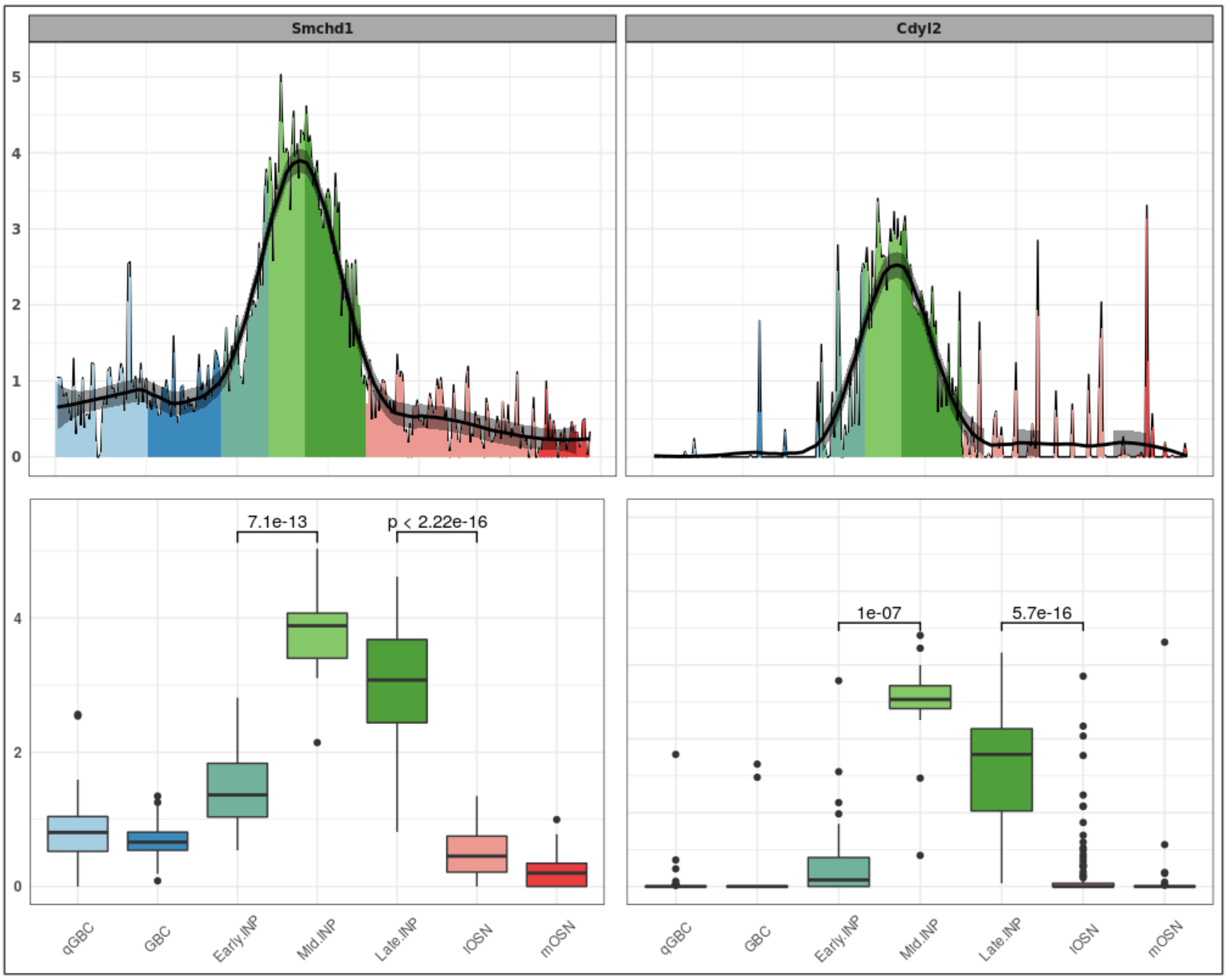

### Pseudotime expression profiles for other factors

**SFigure8: A) pseudotime expression profiles of Kdm1a and factors that play a role in changing the function of Kdm1a. B) Feedback Signal.** Translation of the newly transcribed OR mRNA activates a co-opted arm of the unfolded protein response (Dalton et al., 2013) and induces a feedback signal (Lewcock and Reed, 2004; Serizawa et al., 2005; Shykind et al., 2004). Atf5 and Adcy3 are proposed to be involved in this process where Atf5 promotes Adcy3 expression which in turn downregulates Kdm1a (LSD1) preventing the de-silencing of another OR gene (Dalton et al., 2013; Lyons et al., 2013).

A

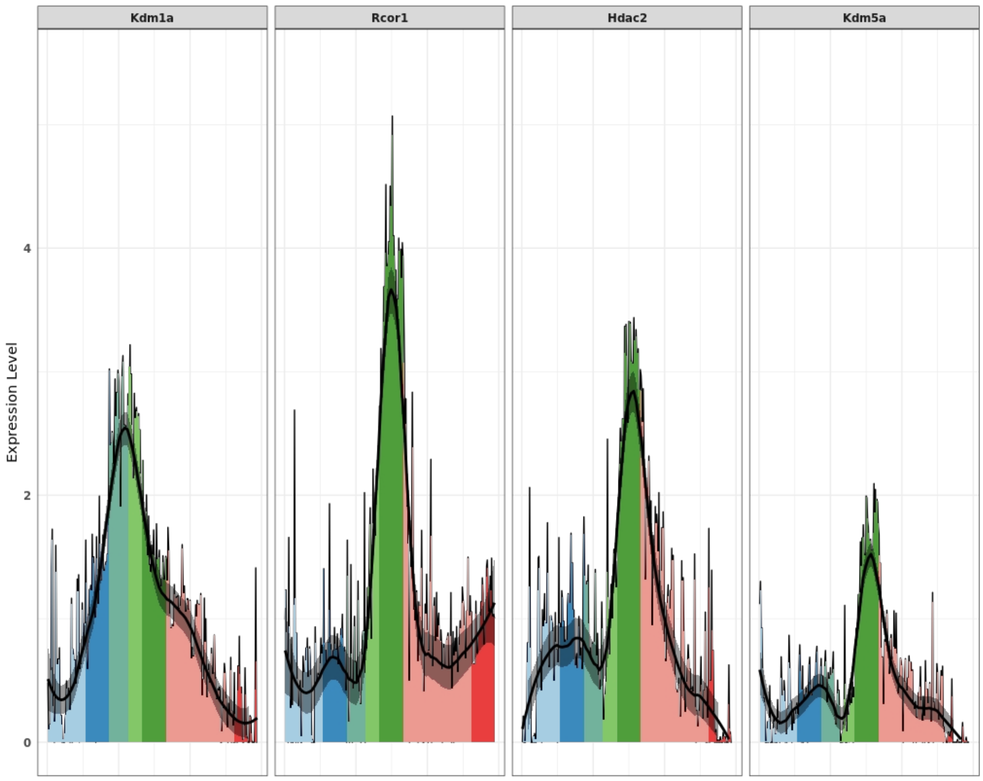


B

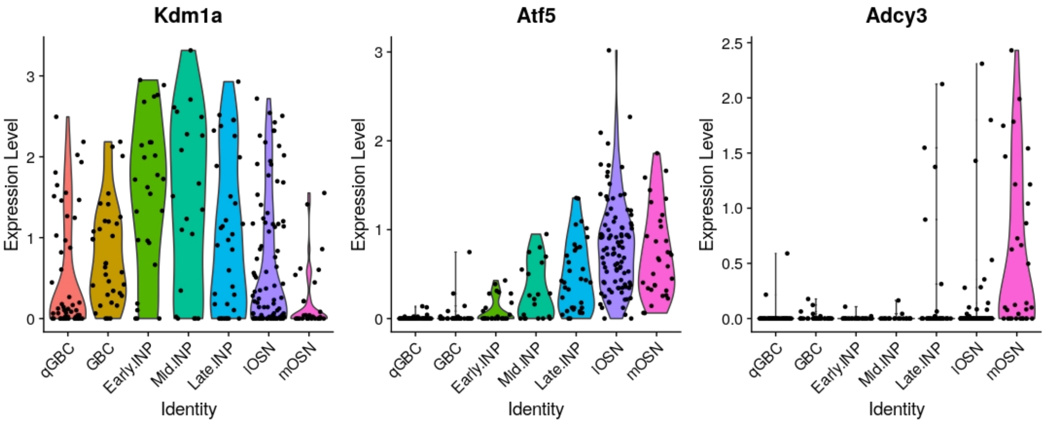

### Pseudotime expression profiles for other factors

**SFigure 9: Other important TFs and chromatin remodelers (CRs) associated with the three phases of OR expression. A)** TFs and co-TFs. **B)** CRs

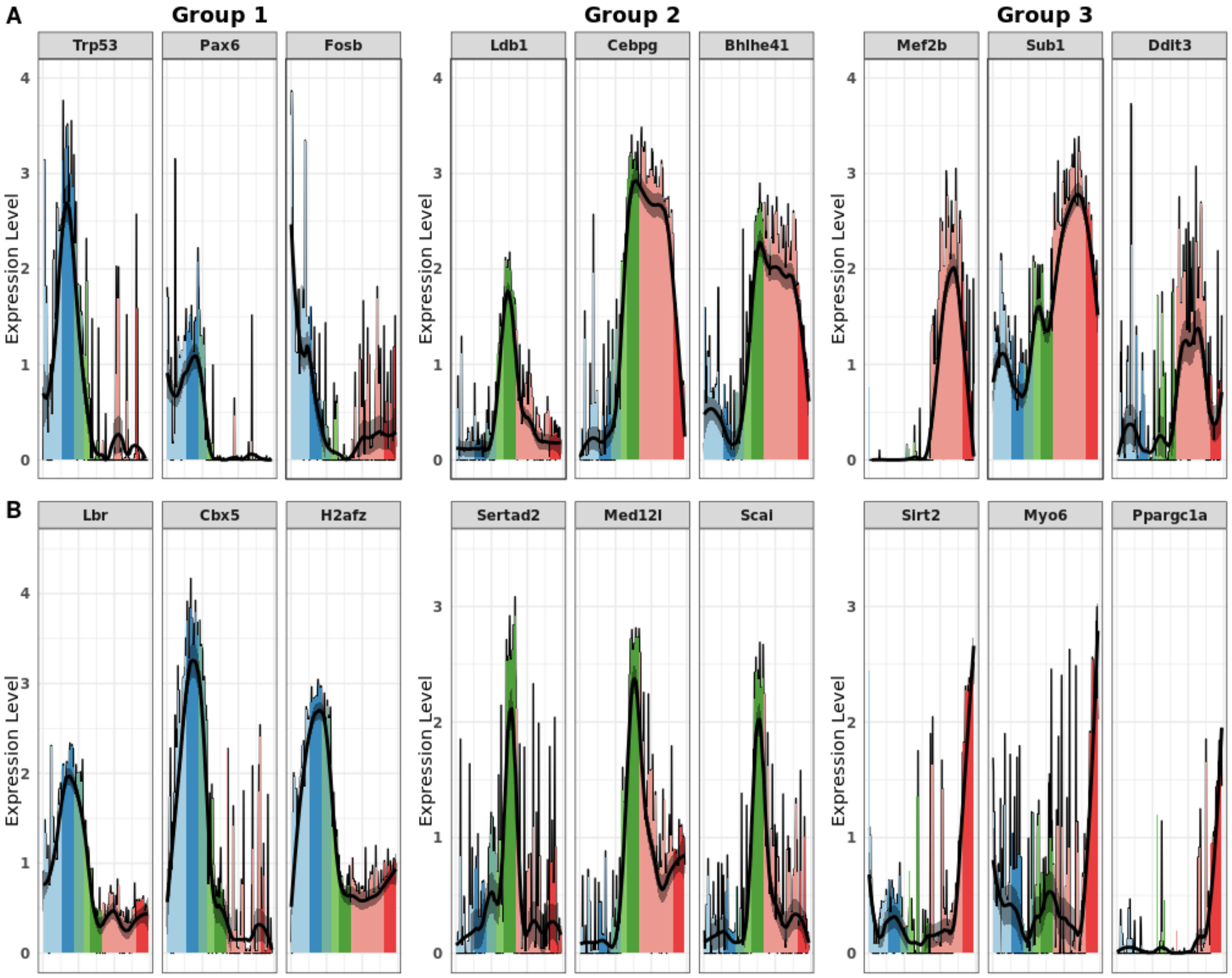

### Coexpression analysis of olfactory receptor genes

**SFigure 10: Coexpression analysis of olfactory receptor genes related to the same cluster.** We performed MCMC sampling to assess whether the observed average number of OR gene clusters co-expressing at least 2 OR genes is in the expected range. We first construct a binary cells x genes matrix indicating whether or not a gene is expressed in a cell. From this matrix, we compute as a test statistic the relative frequency of clusters expressing at least 2 genes among all clusters. The same is done for a uniform random sample of all binary matrices that have the same marginal count frequencies as the original matrix. The random samples are drawn by the curveball algorithm (see SCode 1 below). The figure shows the histogram of the test statistic for this random sample. The red dotted line indicates the actually observed value, corresponding to a (two-sided) p-value of 0.112.

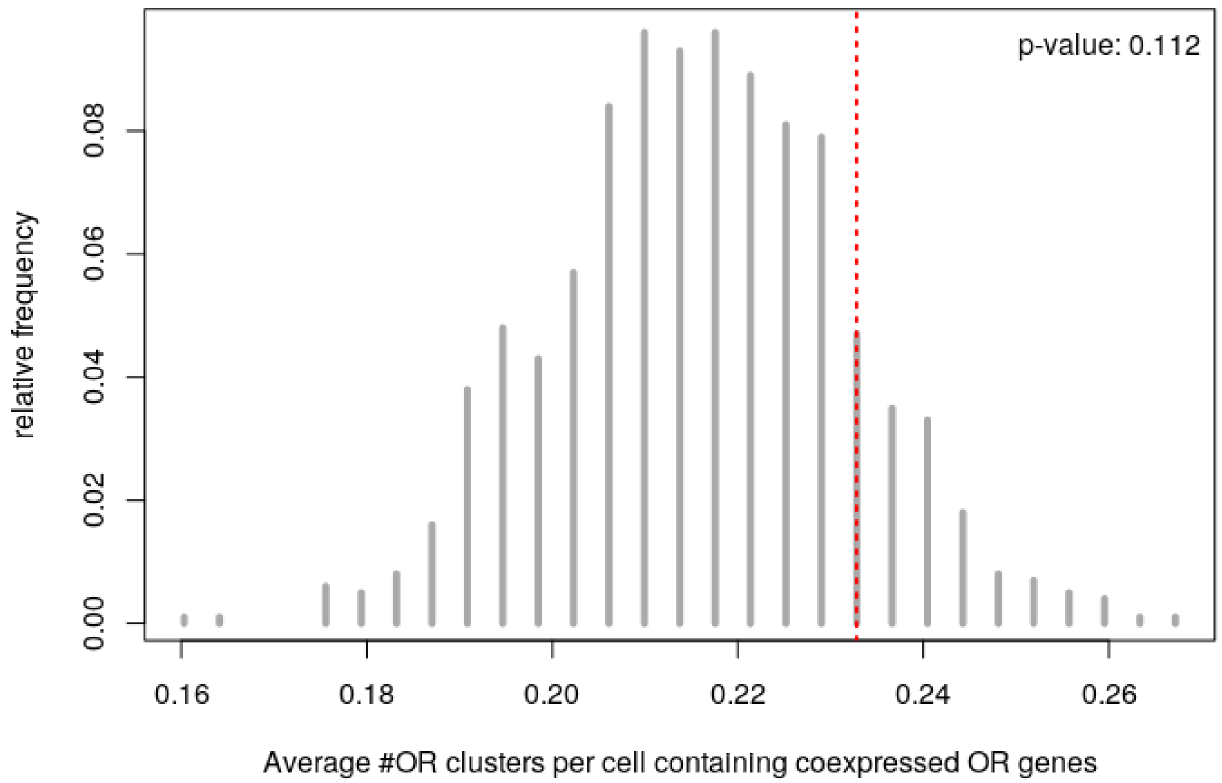

### SCode 1: Coexpression analysis of olfactory receptor genes

We perform MCMC sampling to draw samples uniformly from the set of binary matrices with fixed marginals. We use the 'Curveball' algorithm of Strona et al., Nat.Comm 2014, https://doi.org/10.1038/ncomms5114.

```
# This is the R code provided in Strona et al., Supplementary Software 5

curve_ball<-function(m){

  RC=dim(m); R=RC[1]; C=RC[2]; hp=list()

  for (row in 1:dim(m)[1]) {hp[[row]]=(which(m[row,]==1))}

  l_hp=length(hp)

  for (rep in 1:(5*l_hp)){

    AB=sample(1:l_hp,2); a=hp[[AB[1]]]; b=hp[[AB[2]]]; ab=intersect(a,b)

    l_ab=length(ab); l_a=length(a); l_b=length(b)

    if ((l_ab %in% c(l_a,l_b))==F){

      tot=setdiff(c(a,b),ab); l_tot=length(tot)

      tot=sample(tot, l_tot, replace = FALSE, prob = NULL)

      L=l_a-l_ab; hp[[AB[1]]] = c(ab,tot[1:L])

      hp[[AB[2]]] = c(ab,tot[(L+1):l_tot])}

  }

  rm=matrix(0,R,C)

  for (row in 1:R){rm[row,hp[[row]]]=1}

  rm

}
```

Let expression_mat the original (binarized) genes x cells expression matrix. It contains a 1 whenever the gene is detected in a cell with at least one count. Let assignment_mat the (binary) OR clusters x genes incidence matrix. It contains a 1 at position (c,g) exactly if a gene g is localized in cluster c. Both matrices are available in the Supplementary Materials  (Excel 2).

For each MCMC sample, we record the number of OR genes (co-)expressed in each OR cluster.

```
MCMC_samples = 10^3 # number of MCMC samples we want to draw

MCMC_stats = matrix(0,nrow=nrow(assignment_mat),ncol=MCMC_samples)
```

```
# MCMC_stats contains the number of times an OR cluster had >=2 coexpressed genes
(across all cells), for each OR cluster

mat = expression_mat

# assignment_mat %*% mat is a OR clusters x cells matrix

# containing the #active OR genes (in a given cluster and a given cell)

# MCMC_stats[,j] contains, for each OR cluster the #cells in which

# coexpression  found for this cluster.

MCMC_stats[,1] = rowSums((assignment_mat %*% mat) > 1)

# The original expression matrix is counted as the first sample

cat("Progress: ")

for (j in 2:MCMC_samples){

  mat = curve_ball(mat)

  MCMC_stats[,j] = rowSums((assignment_mat %*% mat) > 1)

  if(j %% (MCMC_samples %/% 25) == 0) cat(".")

}

cat("\n")
```

The test statistic is the relative frequency (in the population of all cells) by which an OR cluster has more than 1 OR gene expressed.

```
cells = ncol(expression_mat)

coexpressed = colSums(MCMC_stats) / cells

# the relative frequency at which we observe coexpressed genes in a cluster

# (across all clusters, per cell), for each MCMC step

original_stats = sum((assignment_mat %*% expression_mat) > 1) / cells

null_distr = table(coexpressed)

location = as.numeric(names(null_distr))

probabilities = as.numeric(null_distr)

probabilities = probabilities/sum(probabilities)

plot(location,probabilities,type="h",lwd=4,col="dark grey",

     xlab="Average #OR clusters per cell containing coexpressed OR genes",
```

```
     ylab="relative frequency")
abline(v=original_stats,col="red",lwd=2,lty=3)


larger = which(location >= original_stats)
pvalue = sum(probabilities[larger])
legend("topright",paste("p-value:",signif(pvalue,digits=3)),bty="n")
```

### Supplementary Data

#### Excel file 1

#### Excel file 2

#### Excel file 3

#### Excel file 4

#### Excel file 5,6

#### Excel file 7
